# DELTA: Training-free and label-free correction of pathology foundation model embeddings through paired counterfactual stain-appearance interventions

**DOI:** 10.64898/2026.11.18.767727

**Authors:** Yoon Ho Choi, Jaewoong Kang, Seung-Ho Shin, Chulho Kim

**Affiliations:** Artificial Intelligence Research Center, Hallym University Chuncheon Sacred Heart Hospital, 77 Sakju-ro, Chuncheon 24253, Republic of Korea; Department of Biomedical Informatics, Hallym University Chuncheon Sacred Heart Hospital, 77 Sakju-ro, Chuncheon 24253, Republic of Korea; Department of Neurology, Hallym University Chuncheon Sacred Heart Hospital, Hallym University College of Medicine, 77 Sakju-ro, Chuncheon 24253, Republic of Korea

**Author notes:** Corresponding authors. E-mail address: Jaewoong Kang, Seung-Ho Shin, and Chulho Kim. First author (Yoon Ho Choi).

**Keywords:** Computational pathology, Pathology foundation models, Stain and scanner variability, Label-free embedding correction, Subspace projection, Domain generalization

## Abstract

Staining and scanning variation impairs pathology foundation model (FM) generalization. We propose DELTA, a training-free, label-free method for post-hoc correction of FM embeddings. DELTA estimates recurrent directions from paired embedding differences between original patches and counterfactual stain-appearance variants preserving spatial structure, then attenuates aligned components through a fixed projection. Estimation from 4% of eligible patches per source dataset required no separately acquired matched images. Operators directly correct stored embeddings. On PathoROB, DELTA improved patch-level representation robustness over raw embeddings in all 20 FMs. Mean classification gains over established correction methods persisted after patch aggregation. DELTA provided additional gains after stain normalization. Whole-slide analyses showed improved in-distribution survival discrimination in TCGA-BRCA and normative cancer screening in a clinical cohort. Operators from TCGA-BRCA reduced embedding variation in the PLISM dataset across 13 staining conditions and 7 scanners without refitting. With projection rank and center matched, DELTA outperformed projections from mismatched pairs in retrieval and tissue-category concordance. Corrected embeddings retained information for spatial gene-expression prediction in the STP-BENCH datasets, with gains for tumor microenvironment markers in selected cohorts.

## Introduction

Hematoxylin and eosin (H&E) histology forms the basis of routine pathological assessment of cellular morphology and tissue architecture. Differences in fixation, tissue processing, section thickness and staining protocols alter color intensity and contrast even among specimens with comparable morphology. Staining practices vary across laboratories and operators, and scanner optics, illumination and color calibration introduce further variation during digitization [1– 4]. In multicenter cohorts, acquisition signatures often coincide with differences in patient and tissue composition. Associations between these signatures and clinical or molecular labels have biased performance estimates and reduced generalization in the prediction of tumor stage, molecular biomarkers and survival [5].

Pathology foundation models (FMs) learn representations from large histopathology collections spanning diverse tissues, institutions and acquisition conditions [6–9]. Embeddings from frozen encoders support tasks ranging from tissue classification and image retrieval to survival modeling. Despite the diversity of pretraining data, analyses across scanners and institutions have identified persistent acquisition-associated differences in FM embeddings and accompanying changes in predictions [10–12]. In the PathoROB benchmark, classification accuracy declined as the association between tissue class and medical center increased in the training data [13]. The decline affected both in-distribution (ID) evaluation on held-out cases from centers represented during training and out-of-distribution (OOD) evaluation on cases from previously unseen centers.

Stain normalization and augmentation address appearance variation in image space. Reinhard, Macenko and Vahadane normalization transform images toward reference color statistics or staining profiles [14–16]. The choice of reference influences the transformed appearance across tissues, and each incoming image requires normalization before embedding extraction. Stain augmentation exposes the encoder to a broader range of colors and stain profiles during training [1]. Its effectiveness depends on how well the simulated transformations represent actual acquisition conditions. Adapting a pretrained encoder through augmentation also requires further training.

Feature-space correction methods target acquisition-associated variation in learned representations. Domain-adversarial neural networks (DANN) optimize representations for task prediction while reducing the accuracy of an adversarial classifier trained to identify acquisition sources [17]. ComBat adjusts the location and scale of embedding coordinates across batches through empirical Bayes estimation [18,19]. Both methods depend on labels identifying acquisition groups. DANN requires these labels for adversarial training and separate annotations for the prediction task. ComBat groups embeddings by batch label and requires samples from a new batch to estimate its adjustment parameters. Depending on the endpoint, task annotations require pathology review, molecular assays or clinical follow-up. Incomplete acquisition records also limit the definition of technical groups. An institutional identifier, for example, groups slides acquired under different staining and scanning conditions. Even with accurate labels, differences between institutions include variation in patient selection and tissue composition. Correction directed at these differences risks attenuating biologically informative variation.

Comparing corresponding tissue regions reduces the influence of differences in tissue content. FEATMAP estimates an affine transformation from paired embeddings of corresponding regions acquired under different conditions [20]. A post-hoc shrinkage method estimates acquisition-induced displacements from matched scanner images and controlled staining perturbations, then attenuates the associated embedding variation [21]. SPECTRA trains low-rank adaptation (LoRA) modules within the encoder on spatially matched tissue images acquired under different staining and scanning conditions. Its contrastive objective requires no task labels [22]. These methods draw on separately acquired image pairs for estimation, calibration or training. Constructing matched collections across scanners and stains entails imaging individual sections on multiple devices and registering serial sections prepared under different staining protocols [4]. Extending coverage to additional institutions requires coordination of specimen preparation, imaging and registration. Even after registration, local differences in tissue content between adjacent sections introduce uncertainty into cross-stain comparisons [22].

We introduce Difference-based Embedding-Level Targeted Adjustment (DELTA), a training-free, label-free method for post-hoc correction of appearance bias in FM embeddings. Paired counterfactual stain-appearance interventions transform a source patch toward the staining appearance of a reference patch while preserving the spatial structure of the source tissue. Differences between the original and transformed embeddings measure the encoder response to these interventions. Recurrent directions across the paired differences define a response subspace. A centered orthogonal projection subtracts the embedding components along this subspace. The operator is estimated separately for each frozen encoder and input preprocessing condition. Estimation requires no additional neural network training, biological or acquisition-source labels, or separately acquired matched images. The operator preserves embedding dimensionality and applies directly to stored or newly extracted embeddings without target-cohort statistics.

We evaluated DELTA across 20 pathology FMs to examine the consequences of correcting individual patch embeddings for downstream prediction. On PathoROB, we assessed acquisition sensitivity in patch representations and evaluated classification under controlled associations between tissue class and acquisition source. Classification gains persisted after patch aggregation. Analyses of whole-slide images (WSIs) extended the evaluation to two distinct clinical tasks. Survival modeling in The Cancer Genome Atlas Breast Invasive Carcinoma (TCGA-BRCA) cohort assessed prognostic information in patient-level representations. Normative screening of gastric biopsies from Hallym University Medical Center (HUMC) assessed cancer detection through local deviations from a non-tumor reference.

To test the relevance of digitally estimated directions to physical acquisition differences, we applied TCGA-BRCA operators to corresponding tissue regions in PLISM [4] without refitting. In comparisons across staining conditions, projections based on principal components or mismatched patch pairs produced distance reductions comparable to those of DELTA at matched rank and projection center. Tissue-category concordance declined with both controls and increased with DELTA. The comparison links tissue discrimination to the directions removed rather than to the magnitude of distance reduction alone. Corrected UNI-v2 embeddings also retained information for spatial gene-expression prediction in STP-BENCH [23]. DELTA separates correction for staining and scanning variation from encoder training, enabling the reuse of stored pathology FM embeddings.

## Results

### Study design

DELTA was evaluated across 20 pretrained pathology FMs to assess representation robustness and downstream prediction after correction of individual patch embeddings (Fig. 1). Patch-level analyses in PathoROB examined acquisition-source sensitivity, tissue discrimination and classification. At the patch-aggregated level, benchmark patches were grouped by slide or patient, and the embeddings within each group were combined into a single representation for classification. WSI-level analyses addressed survival prediction in TCGA-BRCA and normative cancer screening in the HUMC gastric biopsy cohort. Raw encoder embeddings provided the baseline. Reinhard stain normalization, DANN and ComBat served as comparators with distinct requirements for training, labels and evaluation-cohort data (Table 1). Sensitivity to the bank sampling fraction was examined in patch-level and survival analyses (Supplementary Fig. 1).

**Table 1.** Comparison of correction methods.

| **Characteristic** | **Stain normalization** | **DANN** | **ComBat** | **DELTA** |
|---|---|---|---|---|
| Correction mechanism | Fixed-reference color transformation | Learned feature transformation | Batch-wise affine adjustment | Low-rank embedding projection |
| Task labels for fitting | No | Yes | No | No |
| Source or batch labels for fitting | No | Yes | Yes | No |
| Additional feature network training | No | Yes | No | No |
| Image transformation at inference | Yes | No | No | No |
| Target-cohort data required for correction | No | No | Yes | No |
| Direct use of precomputed embeddings | No | Yes | Yes | Yes |

Fitting refers to estimation of the correction itself. Stain normalization refers to the Reinhard method, and DANN denotes domain-adversarial neural networks. ComBat was evaluated transductively, with adjustment parameters estimated from target-cohort embeddings and batch labels.

**Figure 1.**
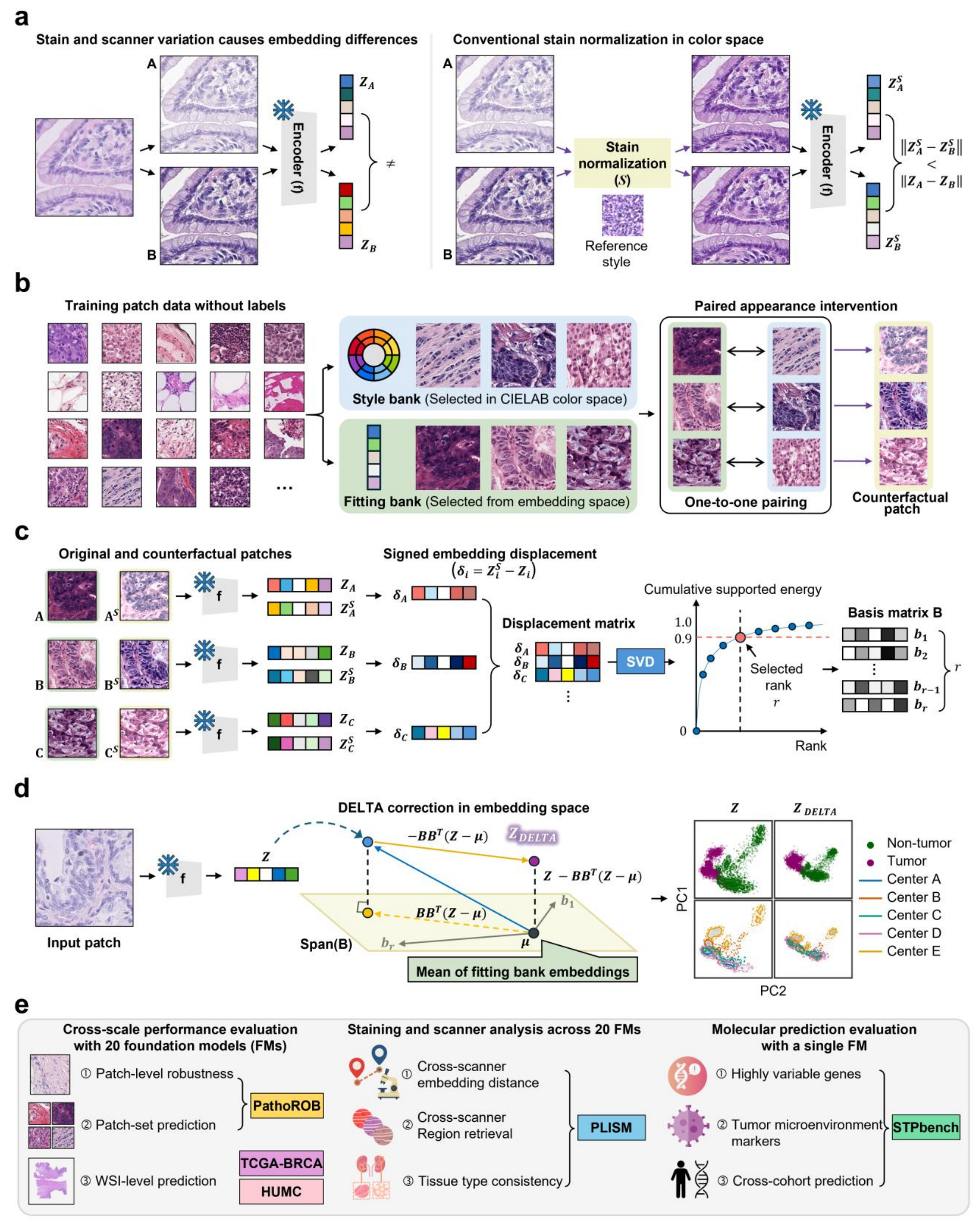
DELTA operator estimation and application. **a** Appearance-associated embedding differences and partial alignment after stain normalization. Distances and embedding maps are schematic. Panel b allocates eligible patches to disjoint Style and Fitting banks and pairs them by retained order. Panel c encodes original patches and their stain-appearance variants, stacks the paired embedding differences and selects a response basis from the supported singular spectrum. Panel d applies the stored center and basis to new embeddings. Panel e summarizes the patch, patch-aggregation, WSI, physical-acquisition and molecular-prediction evaluations. Biological, source and outcome labels are excluded from operator estimation.

Controlled comparisons in PLISM tested the relevance of digitally estimated directions to actual staining and scanner variation. Operators estimated from TCGA-BRCA were applied without refitting and compared with alternative projections matched in rank and center to assess the contribution of paired embedding differences to correction. Spatial gene-expression prediction in STP-BENCH assessed the molecular information retained in corrected UNI-v2 embeddings.

### DELTA improves representation robustness and patch classification

The evaluation covered all four patch datasets in PathoROB [13] and adopted the benchmark’s representation metrics and classification design. Camelyon and Tolkach ESCA provided lymph node and esophageal tissue patches, respectively. TCGA4×4 comprised four cancer classes, and each TCGA2×2 comparison included patches from two cancer classes and two medical centers. RAW and R-RAW denote raw encoder embeddings from original and Reinhard-normalized images, respectively, before embedding correction. The prefix R-identifies Reinhard normalization of the input images across methods. Separate DELTA operators were estimated for each FM in the two input conditions. Figure 2 shows differences in metric scores for DELTA and R-DELTA relative to the indicated comparators.

**Figure 2.**
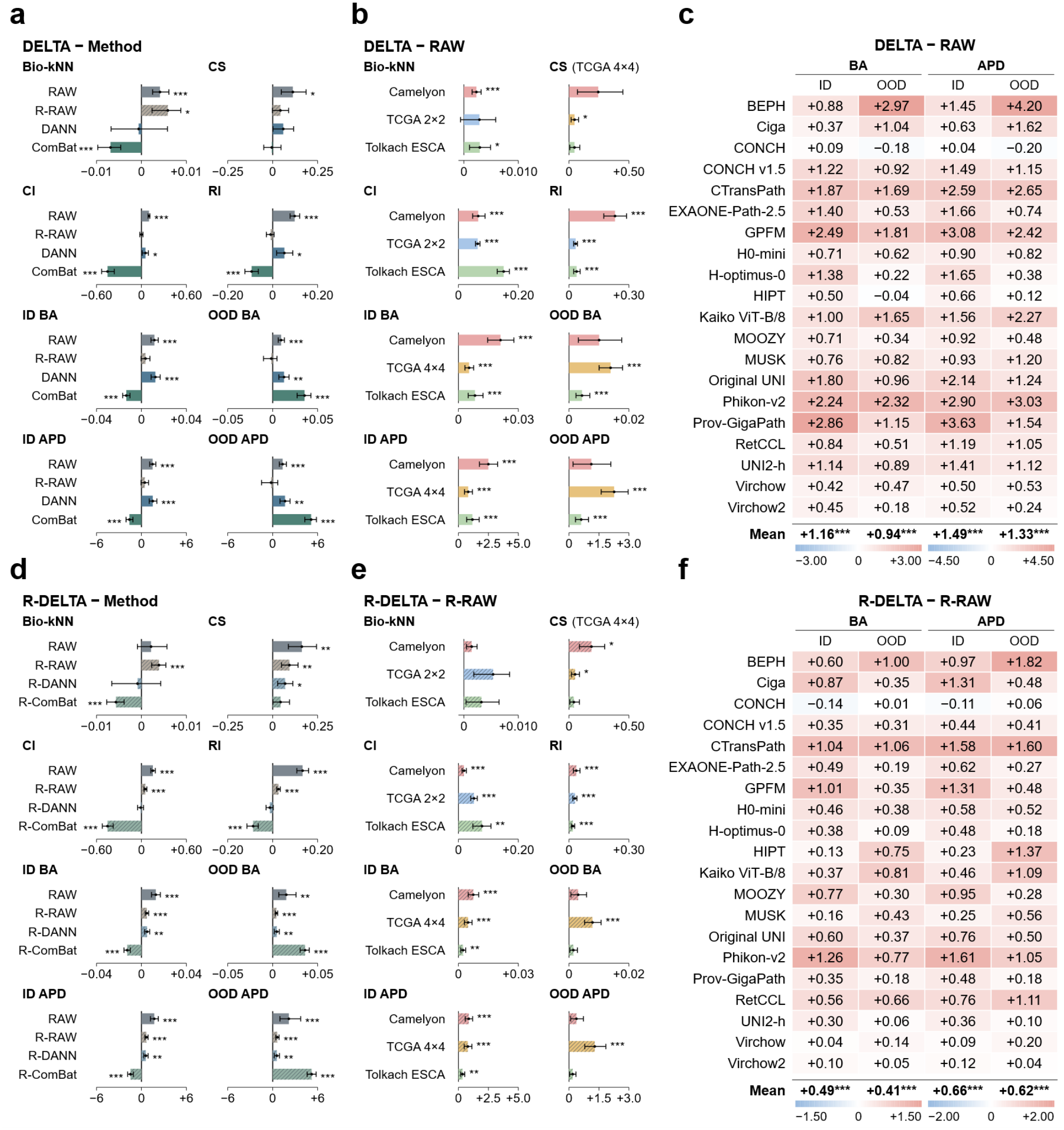
Patch-level robustness of DELTA. Panels **a–c** evaluate DELTA on raw embeddings extracted from original images and **d–f** on embeddings extracted after Reinhard stain normalization. RAW denotes uncorrected embeddings and R denotes Reinhard normalization. **a**,**d**, Mean paired performance differences across 20 backbones. Comparators include RAW, R-RAW, domain adversarial neural networks (DANN) and ComBat, which was evaluated in a transductive setting. **b**,**e**, Dataset-specific differences from RAW and R-RAW, respectively. Pooled representation metrics average Camelyon, TCGA4×4 and Tolkach ESCA. TCGA2×2 supplies dataset-specific neighborhood metrics, and TCGA4×4 supplies clustering and classification. **c**,**f**, Differences in BA and APD for each backbone, comparing DELTA with RAW and R-DELTA with R-RAW, respectively. Positive values favor DELTA throughout. Metrics are biological k-nearest-neighbor performance (Bio-kNN), clustering score (CS), confounder insensitivity (CI), robustness index (RI), balanced accuracy (BA) and average performance drop (APD). ID and OOD denote in-distribution and out-of-distribution evaluation, respectively. BA differences use the accuracy scale in bar plots and percentage points (pp) in heatmaps. APD is calculated relative to a common RAW balanced-condition reference, with differences expressed in pp. In heatmaps, red and blue indicate improvement and deterioration, respectively. Error bars show pointwise 95% confidence intervals from 10,000 bootstrap resamples of 20 paired backbone effects. *P* values were obtained using two-sided exact paired sign-flip tests with Holm correction within comparison families. * *P* < 0.05, ** *P* < 0.01 and *** *P* < 0.001.

The robustness index (RI) contrasts neighbors from other centers sharing the query patch’s tissue category with neighbors from its own center belonging to a different category. For the pooled comparison, results from Camelyon, TCGA4×4 and Tolkach ESCA were averaged within each FM. DELTA increased mean RI from 0.619 with RAW to 0.717, with gains in all 20 FMs. The improvement in mean RI was reproduced after Reinhard stain normalization in the comparison between R-DELTA and R-RAW. Biological nearest-neighbor performance also improved relative to the corresponding raw embeddings in both input conditions (Fig. 2a,d and Supplementary Table 1).

The classification experiments varied the association between tissue category and medical center across training splits. Test patches came from centers represented in classifier training (ID) or from held-out centers (OOD). DELTA and R-DELTA increased mean balanced accuracy (BA) in both domains relative to RAW and R-RAW, respectively. Average performance drop (APD) summarizes the relative change in BA from the common RAW reference under balanced training, with differences in APD reported in percentage points (pp). At held-out centers, APD improved from −3.45% to −2.12% with DELTA (1.33 pp) and from −1.91% to −1.29% with R-DELTA (0.62 pp). Improvements relative to the corresponding raw embeddings remained significant after Holm adjustment for all eight pooled endpoints in both input conditions (Fig. 2a,d).

Compared with Reinhard stain normalization alone (R-RAW), DELTA achieved higher biological nearest-neighbor performance, with the difference remaining significant after Holm adjustment. Under matched preprocessing, DELTA also achieved higher mean BA than DANN in both domains and higher OOD BA than ComBat evaluated transductively (Fig. 2a,d).

The dataset-specific TCGA analyses assessed neighborhood measures on TCGA2×2 and clustering and classification on TCGA4×4. RI and confounder insensitivity improved over the corresponding raw embeddings in Camelyon, TCGA2×2 and Tolkach ESCA in both input conditions. Camelyon showed the largest RI increase over RAW (0.231). Mean BA increased in all three classification datasets in both domains, with the largest gains in Camelyon for ID evaluation and in TCGA4×4 for OOD evaluation (Fig. 2b,e).

At the FM level, DELTA improved both ID BA and APD over RAW in all 20 FMs and both OOD measures in most FMs. After Reinhard normalization, R-DELTA provided additional gains over R-RAW on both OOD classification measures in all 20 FMs (Fig. 2c,f).

### DELTA improves classification from aggregated patch embeddings

The patch-aggregated evaluation extended classification from individual patches to groups of patches from each slide or patient. Within each group, the DELTA-corrected patch embeddings were aggregated by gated attention. Camelyon assessed slide-level tumor detection, and TCGA4×4 assessed patient-level classification of four cancer types. A separate patient-level task within the TCGA4×4 NSCLC subset distinguished lung adenocarcinoma from lung squamous cell carcinoma.

The pooled analysis averaged each metric across the three tasks within each FM, with balanced accuracy (BA) as the primary endpoint. DELTA increased mean BA relative to RAW by 1.75 pp in ID evaluation and 0.73 pp in OOD evaluation. Macro-F1, AUROC and AUPRC also increased relative to RAW. Improvements in all four metrics remained significant after Holm adjustment in both domains (Fig. 3a).

**Figure 3.**
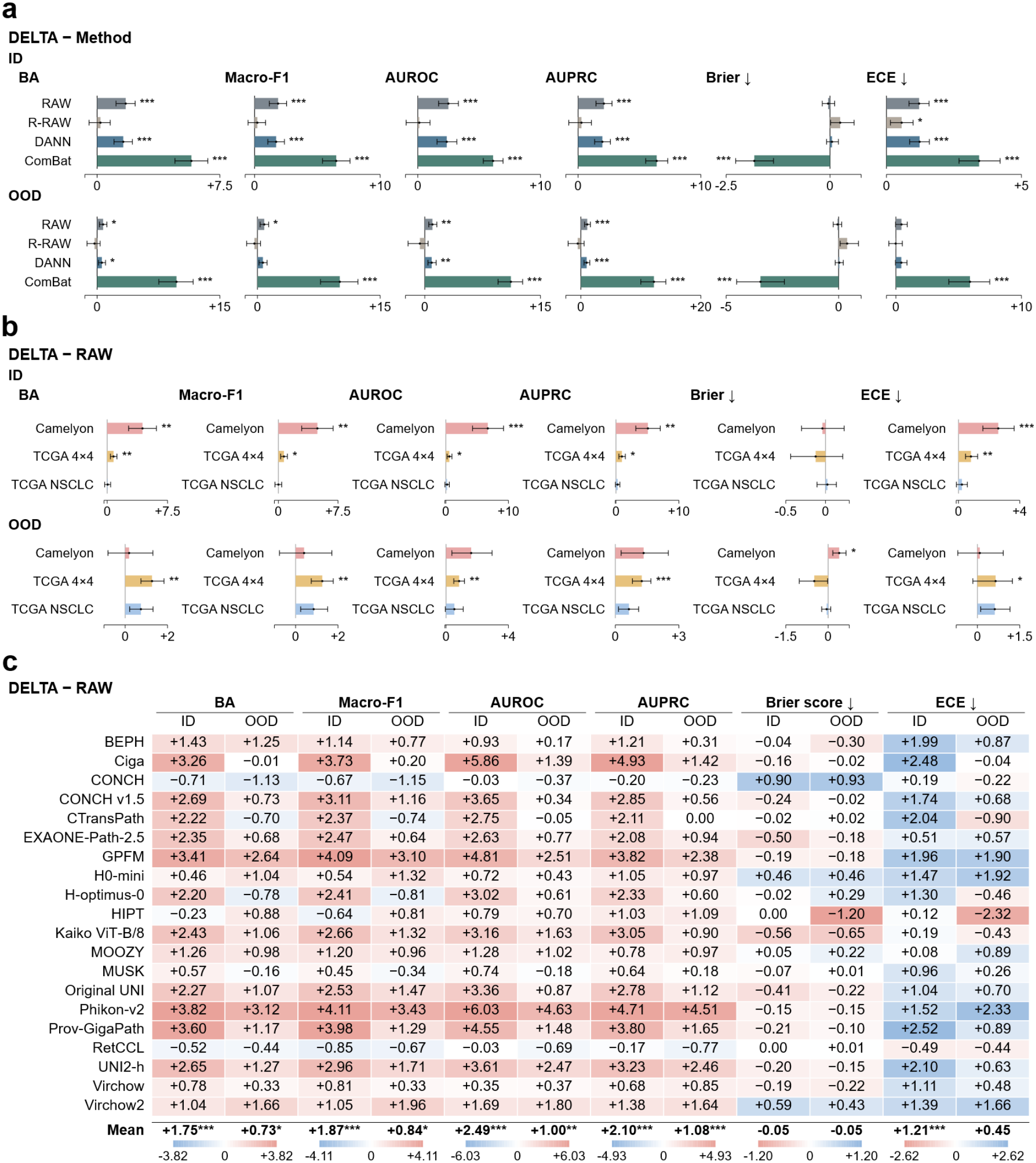
DELTA in patch-aggregated cancer classification. DELTA-corrected patch embeddings underwent gated-attention aggregation for tumor detection in Camelyon, four-class cancer classification in TCGA4×4 and lung adenocarcinoma versus squamous cell carcinoma classification in TCGA NSCLC. Balanced accuracy (BA) was primary. **a**, Mean paired differences across 20 backbones. **b**, Task-specific DELTA-minus-RAW differences. **c**, DELTA versus RAW for individual backbones. Estimates average 100 fixed splits per task. RAW and R-RAW denote uncorrected embeddings from original and Reinhard-normalized images, respectively. ComBat was transductive. Effects are DELTA minus comparator, multiplied by 100. Positive effects favor DELTA for BA, macro-F1, AUROC and AUPRC, and negative effects for Brier score and expected calibration error (ECE). ID and OOD denote in-distribution and out-of-distribution evaluation. Heatmap red and blue indicate improvement and deterioration. Colors reverse for Brier score and ECE, with numerical signs unchanged. Error bars show pointwise 95% confidence intervals from 10,000 bootstrap resamples of paired backbone effects. *P* values are from two-sided exact paired signed-rank tests with Holm correction within comparison families. \**P* < 0.05, \*\**P* < 0.01 and \*\*\**P* < 0.001.

Mean BA was higher with DELTA than with RAW in all three tasks in both domains. The largest ID gain occurred in Camelyon (4.39 pp), and the largest OOD gain occurred in TCGA4×4 (1.27 pp). Both differences remained significant after Holm adjustment (Fig. 3b). In each domain, each discrimination metric favored DELTA over RAW in a majority of FMs. BA improved in 17 of 20 FMs in ID evaluation and 14 of 20 in OOD evaluation (Fig. 3c). Absolute performance by task is reported in Supplementary Data 1.

DELTA also achieved higher mean BA than DANN and ComBat in both domains. Relative to ComBat evaluated transductively, DELTA achieved higher OOD BA in all 20 FMs after averaging across tasks, with a mean difference of 9.70 pp (Fig. 3a and Source Data).

Mean Brier scores were close to those of RAW and lower than those of ComBat in both domains. Among the task-specific comparisons with RAW, only Camelyon in OOD evaluation showed a significant increase in Brier score. In the pooled comparisons, significant adverse differences were confined to expected calibration error (ECE), including an increase of 0.0121 relative to RAW in ID evaluation (Fig. 3a,b).

Mean BA differed little between DELTA and Reinhard stain normalization alone (R-RAW) in either domain (Fig. 3a). After Reinhard normalization, R-DELTA increased mean OOD BA by 0.62 pp over R-RAW. OOD macro-F1 and AUPRC also improved, and all three differences remained significant after Holm adjustment (Supplementary Fig. 2).

### WSI-level survival analysis and normative screening

At the WSI level, survival analysis in TCGA-BRCA assessed the ranking of patients by risk over follow-up (Fig. 4a,c). In the HUMC gastric biopsy cohort, normative screening assessed cancer detection against a reference derived from non-tumor training biopsies (Fig. 4b,d).

**Figure 4.**
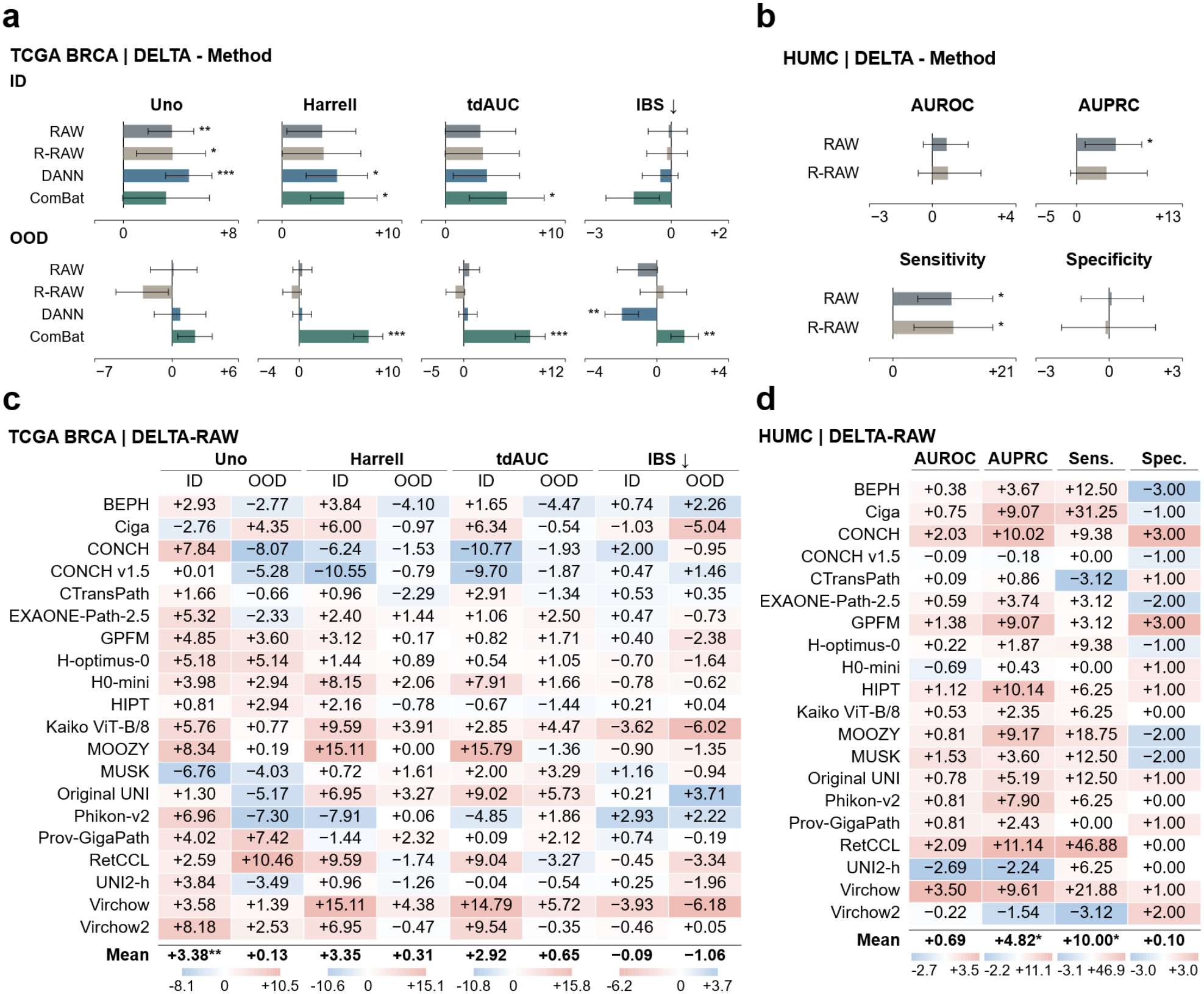
WSI-level survival analysis and normative screening. DELTA was applied to patch embeddings before downstream prediction. **a**,**c**, TCGA-BRCA survival prediction under in-distribution (ID) and held-out-center out-of-distribution (OOD) evaluation. Uno C-index was the primary endpoint, complemented by Harrell C-index, mean time-dependent AUC (tdAUC) and integrated Brier score (IBS). **b**,**d**, HUMC normative cancer screening, assessed using AUROC, AUPRC, sensitivity and specificity from pooled out-of-fold case predictions. Sensitivity and specificity were evaluated at training-derived thresholds. **a**,**b**, Mean paired differences across 20 backbones. TCGA-BRCA comparators include RAW, R-RAW, domain adversarial neural networks (DANN) and transductive ComBat. HUMC comparators include RAW and R-RAW. RAW denotes uncorrected embeddings and R-RAW denotes embeddings extracted after Reinhard stain normalization. **c**,**d**, Corresponding DELTA-minus-RAW differences for individual backbones. All numerical effects are multiplied by 100. Positive differences favor DELTA except for IBS, for which negative differences favor DELTA. Heatmap colors indicate improvement (red) and deterioration (blue), with the color mapping reversed for IBS while numerical signs remain unchanged. Error bars show pointwise 95% confidence intervals from 20,000 bootstrap resamples. TCGA-BRCA intervals resample paired backbone effects. Two-sided exact paired sign-flip P values were Holm-adjusted across four prespecified comparators within each endpoint, evaluation domain and DELTA input branch. HUMC intervals use hierarchical paired resampling of cases within cancer-status strata and backbones. Two-sided centered hierarchical bootstrap P values were Holm-adjusted across five prespecified comparisons within each metric. Heatmap significance symbols refer to mean paired differences and use the same adjusted P values as the corresponding bar plots. * *P* < 0.05, ** *P* < 0.01 and *** *P* < 0.001.

The primary survival endpoint was Uno C-index, which measures agreement between predicted risk rankings and observed survival while accounting for censoring. In ID evaluation, mean Uno C-index increased from 0.6104 with RAW to 0.6442 with DELTA. The mean paired difference remained significant after Holm adjustment, as did the advantages over Reinhard stain normalization alone (R-RAW) and DANN. In OOD evaluation at held-out centers, the mean difference from RAW was close to zero (Fig. 4a and Supplementary Table 2).

The HUMC cohort comprised 132 patients, including 32 with cancer. At thresholds derived from non-tumor training cases, DELTA detected a larger proportion of cancers while flagging a similar proportion of non-tumor cases. Mean sensitivity increased from 75.6% with RAW to 85.6% with DELTA, with mean observed specificity near 94.2% in both conditions. Mean AUPRC also increased, and the gains in sensitivity and AUPRC remained significant after Holm adjustment (Fig. 4b and Supplementary Table 3).

Improvements relative to RAW were observed in most FMs for each task. ID Uno C-index was higher in 18 of 20 FMs in TCGA-BRCA, and screening sensitivity was higher in 15 of 20 FMs in HUMC (Fig. 4c,d).

DELTA also achieved higher screening sensitivity than Reinhard stain normalization alone. The sensitivity improvement over the uncorrected embeddings was reproduced after Reinhard normalization, with R-DELTA increasing mean sensitivity by 8.1 pp over R-RAW at similar observed specificity. Both sensitivity differences from R-RAW remained significant after Holm adjustment (Fig. 4b and Supplementary Fig. 3b,d).

### Controlled validation of the DELTA response subspace in PLISM

Controlled comparisons in PLISM [4] evaluated DELTA across 13 staining conditions and 7 scanners. Operators estimated in TCGA-BRCA were applied to PLISM without refitting. The spatially aligned tissue regions allowed separate comparisons of scanner changes, staining changes and their combination. Scanner comparisons held the tissue section fixed, and comparisons across stains involved registered serial sections (Supplementary Fig. 4).

For operator estimation, the Style bank supplied references spanning variation in staining appearance, and the Fitting bank supplied source patches selected for diversity in the encoder embedding space. Each Fitting patch was digitally transformed toward its assigned Style reference. The response subspace was estimated from the embedding differences between each original Fitting patch and its own counterfactual variant. The unpaired control retained both patch sets but assigned a different original partner to each transformed patch. Projections based on unpaired displacements, Fitting-bank PCA and random orthonormal subspaces were matched to DELTA in rank and center.

Across the three acquisition contrasts, DELTA improved top-1 retrieval of corresponding regions by 1.29–2.36 pp relative to the unpaired control. Tissue-category concordance with the nearest region after exclusion of the corresponding match increased by 1.38–2.06 pp. Both measures also favored DELTA over Fitting-bank PCA and random projections in every contrast. All 18 retrieval and concordance differences against the matched controls remained significant after Holm adjustment (adjusted P < 0.001 for all, Fig. 5a).

**Figure 5.**
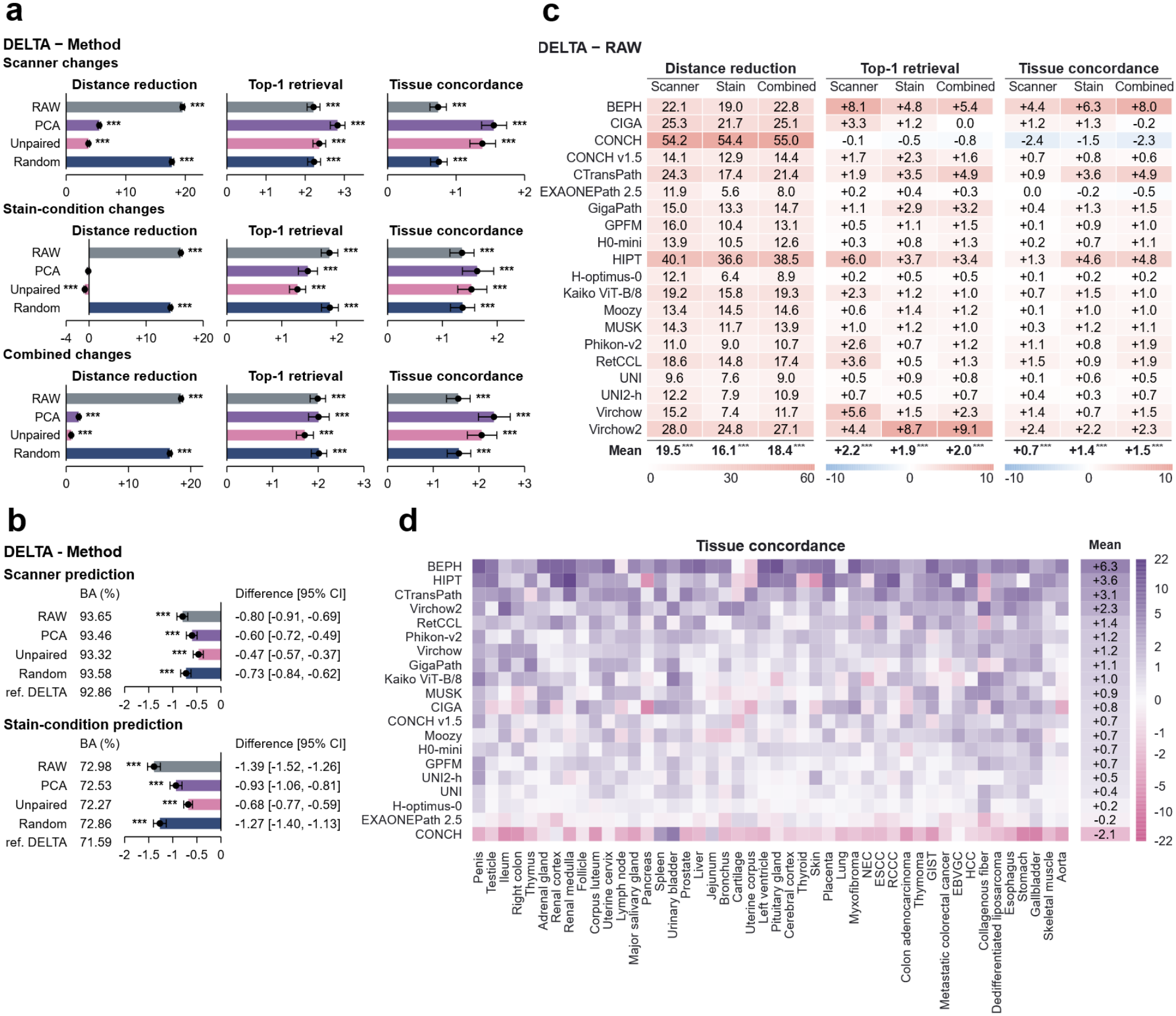
Acquisition-associated variation and tissue retrieval after DELTA correction in PLISM. Frozen TCGA-BRCA-derived operators were evaluated across 20 backbones using 3,417 registered regions from 46 tissue groups, 13 H&E staining conditions and seven scanners. Comparators were uncorrected embeddings (RAW), Fitting-bank PCA, unpaired displacement correction and random projections. Results for random projections were averaged across five orthonormal bases. Correction controls used the DELTA rank and center. **a**, Mean DELTA-minus-comparator differences under scanner, stain-condition and combined changes. Distance reduction measures the percentage reduction in squared Euclidean distance relative to RAW. Top-1 retrieval identifies the corresponding registered region. Tissue concordance measures tissue-label agreement with the nearest gallery region after excluding the corresponding region. Differences are expressed in percentage points (pp). **b**, Scanner and stain-condition prediction using linear ridge-classification probes with five-fold tissue-group-held-out cross-validation. Balanced accuracy (BA) is shown as a percentage, with differences from each comparator in pp. Negative differences indicate lower acquisition predictability under DELTA. **c**, Backbone-specific DELTA-minus-RAW changes. Distance reduction is expressed as a percentage relative to RAW, whereas retrieval and concordance changes are in pp. **d**, Tissue-group-specific concordance changes averaged equally across the three acquisition contrasts. The Mean column averages tissue groups. Purple and pink indicate positive and negative changes. Colors use a shared inverse-hyperbolic-sine mapping with ticks in original percentage-point units. Error bars show pointwise 95% paired percentile bootstrap confidence intervals from 20,000 resamples of the 46 tissue groups, with backbones fixed. Two-sided conditional exact Wilcoxon signed-rank tests used Holm correction across 36 comparisons in **a** and eight in **b**. Mean-row symbols in **c** reuse the corresponding tests in **a**. * *P* < 0.05, ** *P* < 0.01 and *** *P* < 0.001 after correction. Inference is conditional on the observed panel because donor independence is unverified. Probe intervals condition on fitted out-of-fold models. Cross-stain comparisons include serial-section differences.

In the staining comparison, the unpaired projection reduced squared embedding distances slightly more than DELTA, and Fitting-bank PCA reduced them by a similar amount. Tissue-category concordance fell below RAW with both controls and increased by 1.36 pp with DELTA. The advantage over the unpaired control supports the contribution of estimating the response from each original patch and its own counterfactual variant to retaining tissue discrimination after correction (Fig. 5a).

Relative to RAW, mean squared embedding distances decreased by 16.1–19.5% across the three contrasts, with reductions in all 20 FMs. After averaging across FMs and contrasts, tissue-category concordance increased in 45 of 46 tissue groups (Fig. 5c,d and Supplementary Fig. 5). In held-out tissue groups, linear probes also predicted the scanner and staining condition less accurately from DELTA-corrected embeddings than from RAW (Fig. 5b).

### Spatial gene-expression prediction after DELTA correction

Spatial gene-expression prediction assessed the retention of histological information about local molecular variation after DELTA correction. The analysis followed STP-BENCH [23], a benchmark for predicting gene expression at sampled tissue locations from corresponding H&E patches. Under its standard linear-probe protocol, the weights of the default UNI-v2 encoder remained fixed, and separate predictors were fitted to the uncorrected (RAW) and DELTA-corrected embeddings.

Internal cross-validation evaluated held-out sections within each dataset, whereas external transfer evaluated predictors in separate cohorts. Selected comparisons in both settings showed significant increases in Pearson correlation coefficient (PCC) or reductions in mean absolute error (MAE) after Holm adjustment (Fig. 6a). In the NCCHE-Visium lung adenocarcinoma dataset, internal cross-validation yielded a mean PCC for highly expressed and variable genes (HMHVG-200) of 0.0049 below RAW (Holm-adjusted *P* = 0.00021).

**Figure 6.**
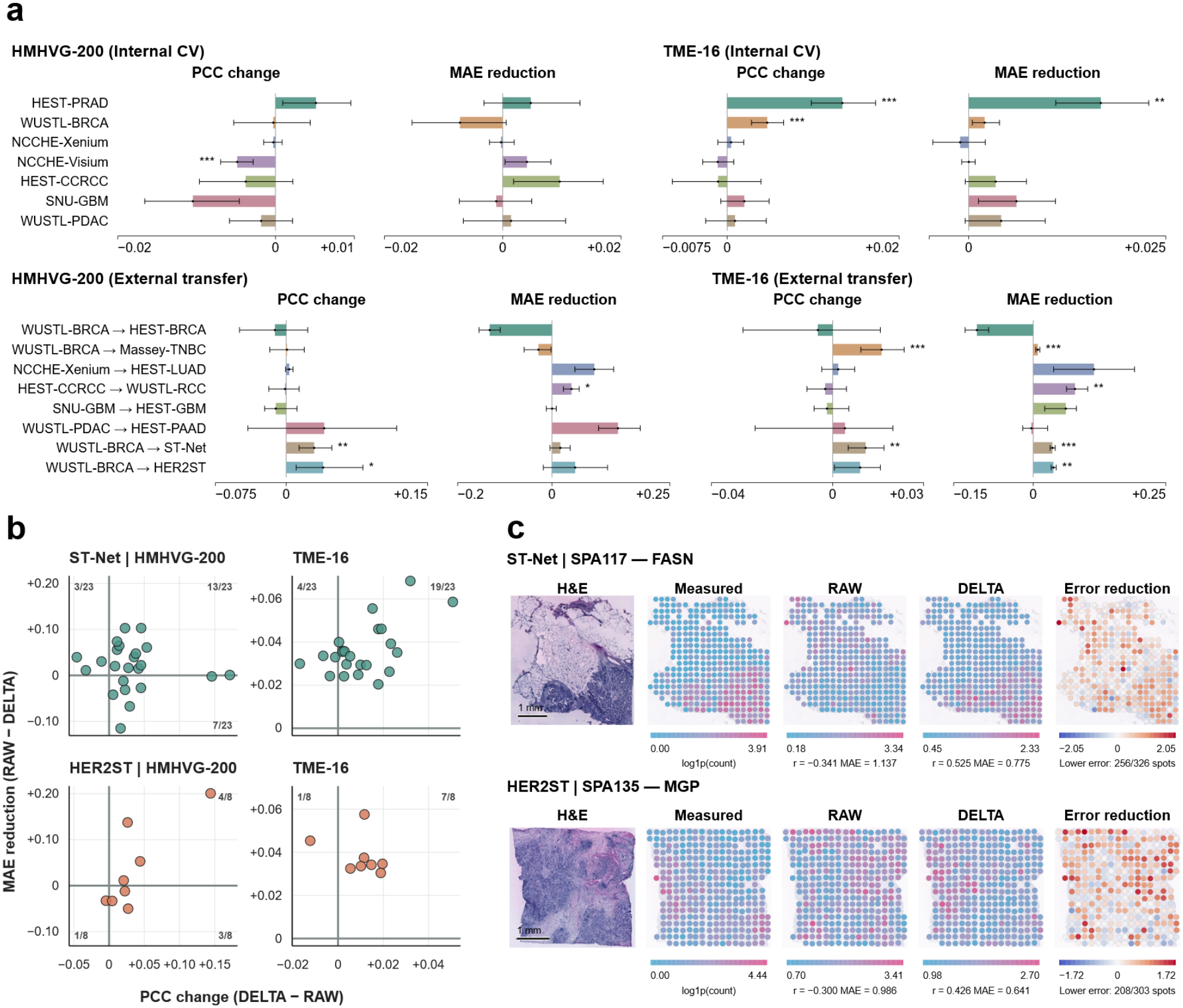
Spatial gene-expression prediction after DELTA correction. Raw and DELTA-corrected UNI-v2 embeddings were evaluated under the STP-BENCH linear-probe protocol. HMHVG-200 and TME-16 denote the trained source gene panels. External evaluation was restricted to genes shared with each target cohort. **a**, Internal cross-validation across seven datasets and 33 predefined folds, and external transfer across eight tasks. Bars show paired changes in Pearson correlation coefficient (PCC), calculated as DELTA minus RAW, and reductions in mean absolute error (MAE), calculated as RAW minus DELTA. Whiskers indicate pointwise 95% paired bootstrap confidence intervals. Original benchmark analyses resampled sections within fixed folds. ST-Net and HER2ST analyses resampled patients while retaining all their sections. Two-sided paired sign-flip tests were followed by Holm correction across tasks within each gene panel and metric. Internal validation, original external transfers and the two additional breast cohorts formed separate comparison families. Asterisks denote adjusted significance levels of * *P* < 0.05, ** *P* < 0.01 and *** *P* < 0.001. **b**, Patient-level PCC changes and MAE reductions in ST-Net and HER2ST. Each point represents one patient, with section-level metrics averaged within patient. ST-Net comprised 23 patients, and HER2ST comprised eight patients. Quadrant annotations indicate patient counts relative to the cohort total. The upper-right quadrant identifies patients with concurrent PCC improvement and MAE reduction. **c**, Spatial predictions for FASN in ST-Net section SPA117 and MGP in HER2ST section SPA135. These illustrative examples were selected for the largest PCC gains with positive MAE reductions across distinct patients and genes. Measured and predicted expression maps display log1p-transformed counts with separate color limits. Error-reduction maps show the difference between RAW and DELTA absolute prediction errors at each spot. Red indicates lower error after correction, and blue indicates higher error. Counts below these maps report spots with lower absolute error under DELTA. Displayed PCC and MAE values were calculated across all spots in each section. Scale bars, 1 mm.

Predictors trained in WUSTL-BRCA were evaluated in two external breast cancer cohorts, ST-Net and HER2ST. For tumor microenvironment markers shared with the training cohort (TME-16), DELTA lowered MAE in every evaluated patient in ST-Net (23 of 23) and HER2ST (8 of 8). PCC also increased in 19 of 23 and 7 of 8 patients, respectively (Fig. 6b). The cohort-level reductions in MAE were 0.0363 in ST-Net and 0.0381 in HER2ST. PCC for HMHVG-200 increased by 0.0297 and 0.0393, respectively. All four differences were significant after Holm adjustment (Fig. 6a).

Across the evaluated sections, mean PCC increased by 0.114 for *FASN* in ST-Net and by 0.152 for *MGP* in HER2ST, with corresponding mean MAE reductions of 0.103 and 0.030 (Supplementary Data 1). The spatial expression maps show one selected section for each gene (Fig. 6c). Supplementary Fig. 6 illustrates both gains and declines, including lower PCC in NCCHE-Visium. The patient-level results support the retention of histological information relevant to spatial gene-expression prediction after DELTA correction.

## Discussion

Institutional contrasts combine acquisition variation with differences in patient and tissue composition [5,13]. Comparing spatially corresponding tissue regions across acquisitions reduces this ambiguity [20–22]. DELTA establishes correspondence digitally by pairing each original patch with its own transformed counterpart. Each paired embedding difference measures the encoder response to a change in staining appearance within a fixed tissue field and provides the basis for subspace estimation.

Application of TCGA-BRCA operators to PLISM [4] without refitting tested the relevance of the estimated subspace to physical acquisition variation. Orthogonal projection contracts distances along the removed directions, so shorter distances alone are insufficient evidence of selective correction. Across staining conditions, projections based on PCA or mismatched pairs reduced distances by comparable amounts but lowered tissue-category concordance relative to RAW. Fixing the number of removed dimensions and the projection center isolates the choice of subspace. The mismatched control retained the original and transformed patch sets but compared different tissue regions, allowing differences in tissue content to enter the displacement estimates. The higher retrieval accuracy and tissue-category concordance with DELTA support pairing each patch with its own variant during subspace estimation.

Retrieval and concordance in PLISM assess the information needed to identify corresponding tissue regions and distinguish tissue categories across acquisitions. STP-BENCH [23] complements this assessment by testing the prediction of spatial gene-expression patterns within tissue sections. The consistent reduction in prediction error for shared tumor microenvironment markers across patients in both external breast cancer cohorts supports the retention of histological information relevant to local expression after correction.

DELTA operates as a fixed correction layer after encoding. The stored operator applies directly to existing embeddings without repeating feature extraction. It also applies to embeddings from new cohorts prepared with the corresponding encoder and input preprocessing, independently of target labels or cohort statistics. Task predictors are fitted and validated on corrected representations before deployment.

Across the 20 FMs, the overall pattern favored DELTA over RAW. Gains were most consistent for patch-level representation robustness, and mean classification gains persisted after patch aggregation. The gains in the main evaluations were obtained with operators estimated from two disjoint banks comprising a combined 4% of eligible source patches (2% per bank). Further improvements after Reinhard stain normalization [14] suggest residual appearance sensitivity after image color alignment. The downstream effects varied in magnitude and direction across FMs, cohorts and metrics. Improved discrimination in ID patch-aggregated classification was accompanied by higher expected calibration error. WSI-level gains depended more on the endpoint and evaluation setting, and survival discrimination changed little at held-out centers. In spatial prediction, the HMHVG-200 correlation decreased in NCCHE-Visium.

This study has several limitations. The 4% sampling fraction was shared across FMs and tasks rather than optimized for each endpoint. Larger banks yielded further gains in some settings (Supplementary Fig. 1). The estimated subspace also depends on the digital stain transformations and tissues represented during estimation. PLISM supported transfer to the staining and scanner conditions examined, while sensitivity to other acquisition factors, such as optical focus and section preparation, remains to be evaluated separately. Spatial gene-expression prediction was evaluated with UNI-v2. The survival and screening findings warrant confirmation in independent multicenter cohorts.

The findings support DELTA as a fixed correction for reducing sensitivity to image appearance in pretrained pathology embeddings. Its operator is estimated directly from existing histology images without labels, additional neural network training or matched images acquired under controlled staining or scanning conditions.

## Methods

### Ethics approval

The HUMC component of this study was approved by the Institutional Review Board of Hallym University Chuncheon Sacred Heart Hospital (IRB No. 2026-05-006). The requirement for informed consent was waived by the Institutional Review Board.

### Study design and datasets

DELTA was evaluated on embeddings from 20 pretrained pathology foundation models (FMs) at the patch, patch-aggregated and whole-slide image (WSI) levels. Complementary analyses examined staining and scanner variation and molecular information retained after embedding correction.

Patch-level analyses followed the PathoROB benchmark [13]. Camelyon, combining CAMELYON16 and CAMELYON17 [24,25], provided normal and tumor lymph node patches. Tolkach ESCA [26] covered six esophageal tissue classes, and TCGA4×4 [27,28] comprised four cancer classes from breast, colon and lung tissues. Representation organization and tissue classification were evaluated across medical centers in these datasets. TCGA2×2 [27,28] provided 94 comparisons across pairs of cancer classes and medical centers for representation robustness assessment.

Patch aggregation through multiple-instance learning [29] assessed classification at the slide and patient levels. Tasks comprised slide-level tumor detection in Camelyon and patient-level four-class cancer classification in TCGA4×4. A separate binary task within the TCGA NSCLC subset of TCGA4×4 distinguished lung adenocarcinoma from lung squamous cell carcinoma, evaluating histological discrimination within lung tissue.

WSI-level survival analysis assessed prognostic information through overall-survival prediction in TCGA-BRCA [30]. This cohort comprised 1,057 WSIs from 996 patients, with survival outcomes obtained from the Genomic Data Commons [31]. Normative screening evaluated gastric cancer detection through local tissue deviations from a non-tumor reference in the Hallym University Medical Center (HUMC) cohort. The HUMC cohort comprised gastric biopsy specimens collected between December 2024 and December 2025 from 132 distinct patients, including 100 non-tumor cases and 32 biopsy-proven cancer cases. The analysis included 117,277 tissue patches from this cohort.

DELTA correction under controlled staining and scanner variation was evaluated with spatially aligned tissue patches from PLISM [4], covering 46 tissue categories, 13 staining conditions and seven scanners. Operators estimated from TCGA-BRCA were applied without refitting.

Retention of molecular information after DELTA correction was evaluated through spatial gene-expression prediction following STP-BENCH [23]. Seven internal datasets comprised 202 sections across 33 predefined folds, and six benchmark external datasets comprised 65 sections. Additional external transfers evaluated ST-Net and HER2ST breast cohorts. Visium and Xenium measurements covered breast, prostate, renal, lung, brain and pancreatic cancers. Raw and DELTA-corrected encoder embeddings from the frozen UNI-v2 encoder were compared under the STP-BENCH linear-probe protocol. Cohort summaries are provided in Supplementary Data 1. Evaluation procedures are detailed in Supplementary Methods S9–S13, S16 and S17.

### Pathology FM encoders

Twenty pretrained pathology FMs were evaluated, spanning convolutional and transformer architectures. Encoder weights remained fixed throughout the evaluations, and embeddings were extracted in evaluation mode. For each encoder, input dimensions, resizing and cropping procedures, interpolation, and RGB channel normalization were held constant across original patches and their stain-transformed counterparts. Embeddings were extracted according to the feature-selection or pooling rule specified for each model. Separate DELTA operators were estimated for each encoder and input branch in the corresponding embedding space. Model architectures, checkpoint identifiers, input preprocessing and embedding extraction settings are detailed in Supplementary Data 1.

### DELTA operator estimation

Staining appearance influences the embeddings generated by pathology foundation models, even for an unchanged tissue region. DELTA characterizes this influence through paired counterfactual stain-appearance interventions, which digitally transform the staining appearance of each tissue patch while preserving its spatial structure. Differences between the original and transformed embeddings quantify the encoder response to each intervention. Recurring directions across these embedding differences define a subspace associated with sensitivity to the applied appearance transformations. DELTA applies a centered orthogonal projection to remove the components of centered embeddings within this subspace. The operator is estimated independently for each frozen encoder and input branch without biological, clinical outcome, or acquisition-source labels.

#### Bank construction and paired counterfactual interventions

The counterfactual transformations of staining appearance are implemented through conventional stain normalization. Each transformation requires a source patch providing the tissue morphology and spatial structure to be retained and a target patch providing the reference staining appearance.

We designate the collections of target and source patches as the Style bank and the Fitting bank, respectively. The Style bank is constructed to represent a range of staining appearances, whereas the Fitting bank is constructed to represent diverse tissue morphologies through selection based on encoder embeddings.

For predictive evaluations, both banks are constructed exclusively from patches in the training dataset. For Style bank construction, staining appearance is characterized by the mean and standard deviation of each CIELAB channel, yielding six color measurements per patch. For Fitting bank construction, encoder embeddings from the corresponding encoder and input branch are reduced to their first 32 principal-component coordinates without whitening. The native encoder embedding coordinates are retained for displacement estimation and projection.

Training patches with a tissue fraction of at least 50% form the eligible population of *N* patches for patch-level, patch-aggregated and WSI-level predictive assessments. Spatial molecular prediction follows the upstream STP-BENCH patch-quality criteria (Supplementary Methods S17). The sampling fraction *p* determines the number of patches per bank as *n* = ⌊*pN*⌋ and was set to 0.02 in this study (Supplementary Methods S15).

Each color measurement and principal-component coordinate is standardized across eligible patches. Patches are then grouped by similarity separately in the color and encoder embedding spaces. Each selected patch is assigned to either the Style bank or the Fitting bank. Both banks comprise *n* patches arranged in selection order (Algorithm 1 and Supplementary Methods S3).

Each source patch *x*_*i*_ in the Fitting bank is paired with the target patch *t*_*i*_ at the corresponding position in the Style bank. Stain normalization transforms the staining appearance of *x*_*i*_ toward the reference provided by *t*_*i*_, producing 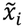. The transformation *T* is implemented with Reinhard stain normalization [14] in this study.

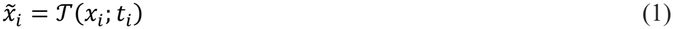

The embedding dimension of the frozen encoder *h* is denoted by *d*. Embeddings of the original and transformed source patches are defined as the *d* × 1 column vectors *z*_*i*_ = *f*(*x*_*i*_) and 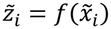, respectively. The displacement *δ*_*i*_ is calculated by subtracting the original embedding from the transformed embedding.

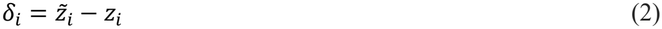

The transposed displacement vectors are arranged as rows of the *n* × *d* displacement matrix *D*. Each row corresponds to one patch pair, and each column corresponds to one encoder embedding coordinate.

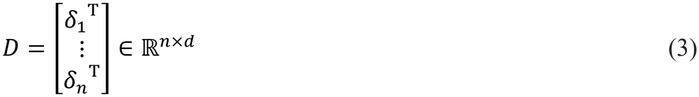

#### Response subspace estimation and embedding projection

Randomized thin singular value decomposition is applied to the displacement matrix *D* to estimate directions recurring across patch displacements. The decomposition approximates the leading *L* components of *D* [32].

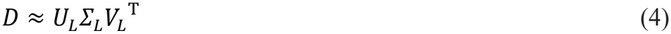

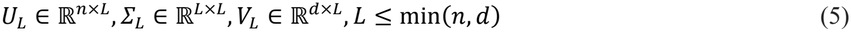

The columns of *U*_*L*_ describe the contribution of each component across patch pairs. The diagonal matrix *Σ*_*L*_ contains the singular values *σ*_*k*_ in descending order. The columns *v*_*k*_ of *V*_*L*_ are defined as mutually orthogonal unit vectors in the encoder embedding space.

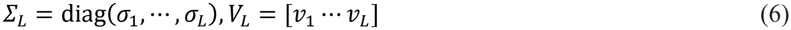

The squared singular value *λ*_*k*_ = *σ*_*k*_^2^ quantifies the contribution of each component to the total squared displacement.

These values are compared with a permutation reference to determine the components included in the projection. One hundred permuted displacement matrices are generated by independently shuffling the row order within each column of *D* . This procedure preserves the distribution of values within each embedding coordinate while disrupting correspondence across coordinates within patch pairs. Squared singular values are calculated for the corresponding components of each permuted matrix. Their 95th percentile defines the reference *q*_*k*_ . The amount by which the observed value *λ*_*k*_ exceeds *q*_*k*_ is defined as *e*_*k*_.

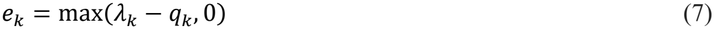

Rank selection considers consecutive leading components satisfying *λ*_*k*_ > *q*_*k*_, starting from the first component. The number of these components is denoted by *r*_*s*_ . The search initially evaluates up to 64 components. If all evaluated components exceed their references, the search width doubles until a component falls at or below its reference or the limit min(*n, d*) is reached. Within this interval, the selected rank *r* is the smallest number of leading components accounting for at least 90% of the summed excess over the permutation references.

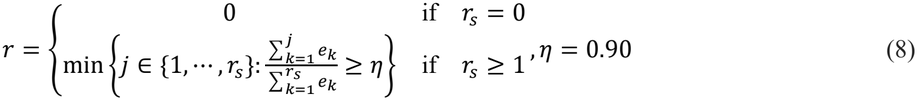

The rank is determined independently for each encoder and input branch. For *r*_*s*_ = 0, the rank is recorded as zero and the identity transformation is applied (Supplementary Methods S6). Survival-specific handling of zero-rank estimates is specified in Supplementary Methods S12.

The selected *r* right singular vectors form the orthonormal basis *B* = [*v*_1_ *⋯ v*_*r*_]. The projection center *µ* is defined as the arithmetic mean of the original Fitting bank embeddings. For a new encoder embedding *z*, the term *BB*^T^(*z* − *µ*) represents the component of the centered embedding within the estimated response subspace. DELTA removes this component through a centered orthogonal projection to obtain the adjusted embedding *z*_*Δ*_.

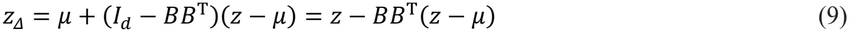

Here, *I*_*d*_ denotes the *d* × *d* identity matrix. This operation preserves the mean of the original Fitting bank embeddings and the output embedding dimension (Supplementary Methods S7).

The estimated *B* and *µ* remain fixed during inference. The operator is applied to new patch embeddings obtained after the preprocessing prescribed for each input branch. For slide- and patient-level tasks, the adjusted patch embeddings are aggregated at the slide or patient level.

### Comparison methods

DELTA was evaluated against raw encoder embeddings and established approaches to stain normalization and embedding adjustment. Stain normalization before embedding extraction was performed with the Reinhard method [14], selected for its previously reported improvements in representation robustness and downstream generalization [13]. For assessments at the patch and patch-aggregated levels and for TCGA-BRCA survival analysis, the normalization reference comprised mean color statistics from 500 TCGA-LUSC patches obtained from 250 patients outside the evaluation cohorts. For HUMC, reference selection was restricted to non-tumor images within each training fold.

Domain-adversarial neural networks (DANN) combined task prediction with adversarial acquisition-source classification through gradient reversal [17]. The learned transformation operated on encoder embeddings and required task and acquisition-source labels during training. All foundation-model encoders remained frozen throughout these comparisons.

ComBat estimated differences in the mean and scale of individual embedding dimensions across medical centers through empirical Bayes estimation [18,19]. Training embeddings were first harmonized across centers to establish a single reference batch. Validation and test partitions were adjusted separately relative to this fixed reference, with medical-center labels defining the batches within each partition.

In summary, stain normalization, DANN and DELTA applied references or transformations fixed before evaluation. ComBat was performed in a transductive manner, with adjustment parameters estimated from the embedding statistics and medical-center labels of each evaluation cohort.

### Patch-level assessment

Patch-level assessment quantified biological organization and acquisition-source sensitivity through biological nearest-neighbor performance, confounder insensitivity, the robustness index and the clustering score defined in PathoROB [13]. Classification was evaluated by balanced accuracy and average performance drop in separate ID and OOD test partitions. ID partitions contained held-out patches from centers represented during classifier training, whereas OOD partitions contained patches from centers excluded from classifier training.

Biological nearest-neighbor performance was the balanced accuracy of predictions obtained by majority vote among neighboring embeddings, with balanced accuracy defined as the mean recall across classes. Neighbor search operated on L2-normalized embeddings and excluded patches from the query slide. Confounder insensitivity quantified the ratio of pooled neighbor counts from other sources to those from the query source. The robustness index contrasted neighbors from other sources sharing the query’s biological class with neighbors from the query source belonging to a different class. The clustering score assessed broader embedding organization through cosine-distance k-means clustering. The silhouette score determined the number of clusters [33], and the adjusted Rand index for source labels was subtracted from the corresponding index for biological labels [34]. Positive scores indicated greater agreement with biological labels than with acquisition-source labels.

Dataset-specific neighborhood analyses covered normal and tumor lymph node patches from Camelyon, six esophageal tissue classes from Tolkach ESCA and 94 comparisons of two cancer classes across two sources from TCGA2×2. Clustering and classification included Camelyon, Tolkach ESCA and the four cancer classes in TCGA4×4. Representation metrics in Fig. 2 were averaged across Camelyon, TCGA4×4 and Tolkach ESCA for each encoder and method. Neighborhood summaries in Source Data were averaged across Camelyon, TCGA2×2 and Tolkach ESCA. Detailed metric definitions, evaluation populations and clustering settings are provided in Supplementary Methods S9.

For supervised classification, predefined splits progressively increased the association between medical center and biological class in training data. Logistic regression was trained separately for each representation, with regularization selected by validation balanced accuracy. DELTA operators were estimated exclusively from training patches. Average performance drop quantified relative changes in balanced accuracy against the fixed RAW balanced-condition reference. Results were averaged within datasets and then equally across datasets within each encoder. The 20 FMs constituted the paired statistical units. Classification partitions, regularization settings and performance-drop calculations are detailed in Supplementary Methods S10.

### Patch-aggregated assessment

Patch-aggregated assessment evaluated slide- and patient-level classification by aggregating encoder embeddings from the Camelyon and TCGA4×4 patch collections examined at the patch level. Balanced accuracy was the primary endpoint, complemented by macro F1, AUROC and AUPRC. Brier score [35] quantified error in predicted probabilities, and expected calibration error [36] assessed agreement between predicted confidence and observed accuracy.

Three classification tasks were evaluated. Camelyon comprised 49 normal and 48 tumor slides for binary classification. TCGA4×4 comprised 272 patients, with 68 per cancer class, for four-class classification. The TCGA NSCLC subset provided 68 patients with lung adenocarcinoma and 68 with lung squamous cell carcinoma for binary classification.

All released patches from each slide or patient were retained. DELTA correction preceded aggregation by a gated-attention multiple-instance learning model [29], with architecture, optimization settings and validation-based model selection held constant across representation branches.

Each task was evaluated over 100 predefined splits spanning different associations between medical center and biological class, with separate ID and OOD test partitions. All patches from each slide and all slides linked by an available patient identifier remained within a single partition. DELTA operators were estimated exclusively from training patches. Grouping rules, partition construction, model specifications and endpoint calculations are detailed in Supplementary Methods S11.

### WSI-level assessment

WSI-level assessment examined the retention of prognostic information and sensitivity to localized tissue abnormalities after DELTA correction. Survival analysis evaluated overall-survival prediction from tissue embeddings averaged at the patient level. Normative screening evaluated gastric cancer detection through local embedding deviations from a non-tumor reference.

#### Survival analysis

Uno C-index [37] was the primary endpoint. Harrell C-index [38] and mean time-dependent AUC [39] provided complementary measures of survival discrimination, and integrated Brier score [40] quantified error in predicted survival probabilities.

The 996 TCGA-BRCA patients were assigned to training (381), validation (84), ID test (80) and OOD test (451) partitions. Corrected embeddings were averaged across all eligible tiles from all slides belonging to each patient. A neural Cox model [41] was trained through Efron partial-likelihood optimization [42], with checkpoint selection by validation Uno C-index. Patient risk predictions were averaged across three optimization seeds before endpoint calculation. Standardization parameters, the censoring distribution and the Breslow baseline hazard [43] were estimated from training patients and remained fixed for ID and OOD evaluation. Event counts, model specifications and survival-metric calculations are detailed in Supplementary Methods S12.

#### Normative screening

AUROC and AUPRC quantified discrimination across case scores. Sensitivity and specificity assessed detection at a fixed threshold. The HUMC analysis included 100 non-tumor gastric biopsy cases and 32 biopsy-proven cancer cases. Within each of five case-level folds, DELTA operators, normative references and thresholds were estimated exclusively from training non-tumor images.

Each patch received an anomaly score defined as its mean cosine distance to the 20 nearest training non-tumor embeddings. Case scores averaged the highest 1% of patch scores, including at least five patches. Each representation branch applied a threshold fixed at the 95th percentile of its leave-one-case-out training non-tumor case scores. Case sampling, fold construction and supplementary operating-point analyses are detailed in Supplementary Methods S13.

### Assessment of staining and scanner variation

DELTA correction was evaluated through separate controlled comparisons across staining conditions and scanners. Spatially aligned image patches provided by PLISM [4] were compared with one acquisition factor held constant while the other differed between images. Scanner comparisons involved an identical physical section, whereas staining comparisons involved serial sections. The H&E condition with Mayer hematoxylin, designated MY, served as the fixed staining reference and provided 3,417 corresponding tissue regions across seven scanners.

Evaluation covered original-input embeddings from 20 FMs. DELTA and comparison operators were estimated from TCGA-BRCA and remained fixed throughout PLISM evaluation. To assess the contribution of paired counterfactual stain-appearance interventions to subspace estimation in DELTA, the orthonormal basis *B* was replaced with a Fitting-bank PCA basis, an unpaired displacement basis or random orthonormal bases. Each alternative operator retained the selected rank and projection center for the corresponding encoder and applied centered orthogonal projection.

The Fitting-bank PCA basis comprised the leading principal components of centered original Fitting-bank embeddings, assessing removal of their dominant variation. The unpaired displacement basis was estimated after permuting the original partners of transformed embeddings with no original pair retained. This control assessed the contribution of correspondence between each source patch and its transformed counterpart. The random orthonormal bases were generated from Gaussian random vectors and assessed removal of arbitrary subspaces of the selected rank.

Squared embedding-distance reduction quantified the fraction of the original squared distance removed by projection between corresponding patches. For tissue region *i*, scanners *a* and *b*, encoder *e*, and correction method *m*, this quantity was defined as

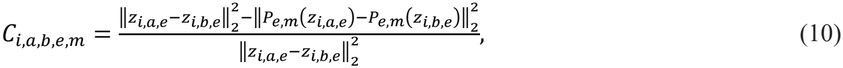

where *z*_*i,a,e*_ denotes the raw encoder embedding of the patch representing region *i* imaged by scanner *a*, and *P*_*e,m*_ denotes the frozen projection operator. This measure also applied to comparisons across staining conditions and to comparisons differing in both acquisition factors.

Matched-field retrieval assessed identification of the corresponding patch across acquisition conditions. Tissue-category concordance measured agreement in tissue category with the nearest neighboring patch after exclusion of the corresponding patch (Supplementary Fig. 5). MY scanner results were summarized descriptively (Supplementary Table 4). Scanner and staining-condition predictability was assessed with linear ridge-classification probes on held-out tissue groups. Inference retained 46 paired tissue groups with the 20-encoder panel fixed. Basis generation, retrieval endpoints, aggregation and statistical procedures are detailed in Supplementary Methods S16.

### Spatial molecular prediction

Spatial gene-expression prediction on datasets from STP-BENCH [23] assessed the retention of molecular information after DELTA correction. Pearson correlation quantified linear correspondence between predicted and measured expression across tissue locations, and mean absolute error quantified prediction error on the transformed expression scale (Supplementary Fig. 6). Analyses covered 200 high-mean highly variable genes (HMHVG) and 16 tumor-microenvironment markers across internal and external Visium and Xenium datasets (Supplementary Data 1).

Under the benchmark linear-probe protocol, separate linear predictors were trained on raw encoder embeddings and DELTA-corrected embeddings from the frozen UNI-v2 encoder, with training folds and optimization settings held constant. DELTA operators were estimated exclusively from source-training images without gene-expression information and remained fixed during validation and external transfer. External evaluation applied the operator and trained predictor from source fold 0. Metrics were calculated for each gene across tissue locations and averaged over evaluable genes within each section. Preprocessing, predictor implementation, aggregation and statistical procedures are detailed in Supplementary Methods S17.

### Statistical analysis

The 20 FMs were treated as paired statistical units in patch-level and patch-aggregated assessments and WSI-level survival analysis. Patch-level results were aggregated across datasets within each encoder. Patch-aggregated results were averaged across 100 splits per task and equally across the three tasks. Paired bootstrap confidence intervals [44] were estimated from 10,000 resamples for patch-level and patch-aggregated assessments and 20,000 for WSI-level survival analysis. Patch-aggregated intervals were pointwise 95% intervals. Model-panel inference was conditional on the observed cohorts, with shared model ancestry potentially limiting encoder independence.

Patch-level contrasts were evaluated with two-sided exact sign-flip tests. Holm adjustment [45] was applied across seven comparators within each representation metric and DELTA input branch, and across six comparators and the ID/OOD domains within each classification metric. Dataset-specific analyses were adjusted separately. Patch-aggregated contrasts employed two-sided exact Wilcoxon signed-rank tests [46], with Holm adjustment across four comparators within each input branch, endpoint and domain. Analyses of individual tasks were adjusted across the four comparators and three tasks. ComBat was included in both assessment levels. Comparator assignments and adjustment procedures are specified in Supplementary Methods S14.

For WSI-level survival analysis, endpoints were calculated from patient risk predictions averaged across optimization seeds. Two-sided exact sign-flip tests evaluated prespecified contrasts, with Holm adjustment across four comparators within each endpoint, evaluation domain and DELTA input branch. Common-control maxT adjustment [47] provided a multiplicity sensitivity analysis.

HUMC screening inference comprised 20,000 hierarchical paired-bootstrap replicates. Non-tumor and cancer cases were resampled separately with shared weights across methods and encoders, followed by resampling of the 20 FMs. Predictions and fold-specific thresholds remained fixed. Two-sided P values from the centered hierarchical difference distribution were Holm-adjusted across five comparisons per endpoint. Resampling procedures and secondary analyses are detailed in Supplementary Methods S14.

Statistical inference for staining and scanner variation assessed distance reduction, retrieval and tissue-category concordance in PLISM. The 46 tissue groups formed the paired statistical units, with the 20 FMs held fixed. Pointwise 95% percentile confidence intervals were calculated from 20,000 paired bootstrap resamples of tissue groups. Two-sided exact Wilcoxon signed-rank tests were Holm-adjusted across 36 comparisons. Scanner and staining-condition prediction was assessed separately, with Holm adjustment across eight comparisons. Confidence intervals for these prediction analyses were conditional on the fitted out-of-fold classifiers. The MY scanner analysis remained descriptive. Acquisition-specific aggregation and inference are detailed in Supplementary Methods S16.

Spatial molecular prediction treated tissue sections within fixed folds as paired sampling units in the original benchmark analyses. ST-Net and HER2ST analyses treated patients as paired sampling units, with all sections from each patient retained during resampling. Paired bootstrap intervals and sign-flip tests were calculated per dataset or transfer, gene panel and metric. Holm adjustment covered tasks separately within the internal analyses, the original external transfers and the additional ST-Net and HER2ST transfers for each gene panel and metric (Supplementary Methods S17).

Missing results were not imputed.

### Implementation details

Experiments were implemented in Python with PyTorch and torchvision. Numerical and statistical analyses were performed with NumPy, scikit-learn and SciPy. GPU computations were performed on NVIDIA GPUs with CUDA acceleration. Software environments and hardware are detailed in Supplementary Methods S18 and Supplementary Table 5.

## Data availability

The PathoROB patch datasets are available through the PathoROB collection on Hugging Face (https://huggingface.co/collections/bifold-pathomics/pathorob). TCGA-BRCA whole-slide images and clinical data are available through the National Cancer Institute Genomic Data Commons Data Portal (https://portal.gdc.cancer.gov/projects/TCGA-BRCA). Spatially aligned PLISM patches are available from Figshare+ (https://doi.org/10.25452/figshare.plus.23614422.v1). The original STP-BENCH analyses [23] used preprocessed data from Hugging Face revision 76510407, subject to the repository’s access conditions. The Massey-TNBC source data are available from NCBI GEO under accession GSE210616. External transfer used fold 0 of the WUSTL-BRCA source partitions and all 43 sections listed in the Massey-TNBC test manifest, both from STP-BENCH commit dd476415. The original studies describe the datasets and access conditions for ST-Net [48] and HER2ST [49]. HUMC data are restricted to institutional access and are not available to external investigators.

## Code availability

Code and frozen configurations required to reproduce DELTA estimation and evaluation will be released at https://github.com/YOONHO-CHOI/DELTA before publication. The release will include representation contracts, bank manifests, pairing records, equal-pair weighting contract, rank-selection outputs, and analysis scripts.

## Acknowledgments

This research was supported by Hallym University Medical Center Research Fund.

## Author contributions

Y.H.C., Conceptualization, Resources, Methodology, Software, Formal analysis, Investigation, Validation, Visualization, Writing - original draft, Writing - review & editing, Project administration; J.K., Methodology, Writing - review & editing, Supervision; S.-H.S., Validation, Writing - review & editing, Supervision; C.K., Writing - review & editing, Supervision.

