## Supplementary Information for "DELTA: Training-free and label-free correction of pathology foundation model embeddings through paired counterfactual stain-appearance interventions"

### Supplementary Methods

#### S1. Scope of operator estimation

Each operator is estimated for one frozen encoder and one input branch. The original-input branch encodes the original images. The Reinhard-normalized branch encodes images normalized to the reference defined in S8. Bank construction and operator estimation are repeated independently in the resulting embedding spaces. Patch-level, patch-aggregated and WSI-level predictive analyses restrict estimation to training patches with tissue fraction of at least 0.50. STP-BENCH [1] eligibility follows its upstream patch-quality criteria without an additional tissue-fraction threshold. Descriptive PathoROB [2] representation analysis estimates operators from the eligible benchmark collections. Biological class, outcome, institution, scanner, data source and task labels are excluded from bank construction, pairing, displacement estimation, rank selection and projection. Predictive validation and test embeddings are also excluded. Supplementary Data 1 distinguishes operator-estimation populations from evaluation material.

The frozen encoder output defines the native representation. DELTA preserves its dimension, allowing existing downstream architectures to be fitted on adjusted embeddings without changing their input interface. Operator estimation includes paired stain-appearance transformations. Inference applies the stored projection to the encoder output, followed by the downstream model fitted on adjusted training embeddings. This correction requires neither new paired image transformations nor statistics from the incoming cohort. The Reinhard-normalized input branch retains its specified image normalization before encoding.

#### S2. Core notation

The notation table below summarizes recurring core symbols. Symbols used only once are defined at first use. All vector and matrix dimensions follow the column-vector convention stated in Supplementary Methods S5.

| Symbol | Definition |
|---|---|
| $\mathcal{E}$ | Eligible patch population for operator estimation |
| $N$ | Number of patches in the eligible population |
| $p$ | Fraction of the eligible population assigned to each bank |
| $\mathcal{S}$ | Ordered Style-bank sequence representing stain appearance |
| $\mathcal{F}$ | Ordered Fitting-bank sequence representing encoder-specific embedding diversity |
| $n$ | Number of patches in each bank and number of one-to-one pairs |
| $x_i$ | Fitting patch in pair i |
| $t_i$ | Style-reference patch in pair i |
| $\tilde{x}_i$ | Counterfactual patch formed by applying the paired Style appearance to the Fitting patch |
| $f$ | Fixed pathology foundation-model encoder |
| $z_i$ | Source embedding of the Fitting patch |
| $\tilde{z}_i$ | Embedding of the paired counterfactual patch |
| $\delta_i$ | Paired Euclidean embedding displacement |
| $w_i$ | Equal normalized bookkeeping weight, fixed as the reciprocal of the retained pair count |
| $D$ | Equal-contribution n by d displacement matrix with one transposed column-vector displacement per row |
| $K$ | Number of strata in each color or embedding coordinate space, defined in Eq. (S6) |
| $L$ | Number of leading singular components evaluated at the current spectral width |
| $U_L$ | n by L matrix of the leading left singular vectors of D |
| ${D_0}^{(b)}$ | n by d displacement matrix after independent coordinate-wise permutations in replicate b |
| $\pi_j^{(b)}$ | Independent permutation of the n pair indices for feature coordinate j in replicate b |
| $\sigma_k$ | Observed singular-value amplitude of component k |

| Symbol | Definition |
|---|---|
| $\sigma_{0,k}^{(b)}$ | Singular-value amplitude of component k in permutation replicate b |
| $\lambda_k$ | Squared observed singular value of component k |
| $\lambda_{0,k}^{(b)}$ | Squared singular value of component k in permutation replicate b |
| $q_k$ | Empirical 95th percentile of the 100 permuted squared singular values for component k |
| $e_k$ | Positive excess of the observed squared singular value above its permutation reference |
| $\tilde{e}_k$ | Excess retained within the leading interval and set to zero outside it |
| $r_s$ | Endpoint of the supported uninterrupted leading interval |
| $r$ | Selected response-subspace rank |
| $B$ | Orthonormal response basis with r columns |
| $\mu$ | Unweighted arithmetic mean of the selected source Fitting embeddings |
| $z_\Delta$ | DELTA-adjusted embedding |
| $\mathcal{X}_{\text{train}}$ | Patch set in the training split |
| $\rho(u)$ | Tissue fraction of patch u |
| $d$ | Native embedding dimension |
| $\mathcal{P}$ | Ordered set of one-to-one Fitting–Style pairs |
| $\mathcal{T}$ | Deterministic stain-appearance transformation used to create a counterfactual patch |
| $Q_{0.95}$ | Empirical component-wise 95th-percentile operator |
| $\eta$ | Required fraction of the summed excess within the leading interval, fixed at 0.90 |
| $z$ | New d by 1 column-vector embedding to be adjusted |
| $I_d$ | d by d identity matrix |
| $I_r$ | r by r identity matrix |
| $B^{*(s)}$ | d by r response basis from bootstrap replicate s, defined when r is at least one |

**S3. Deterministic construction of disjoint Style and Fitting banks**

Let $f$ denote the fixed encoder. For predictive evaluations, the training patch set $\mathcal{X}_{\text{train}}$ is screened by the tissue-fraction function $\rho(u)$. The encoder embedding dimension is denoted by $d$. Equation (S1) defines the eligible population $\mathcal{E}$ and its size $N$. The Style bank represents variation in staining appearance, and the Fitting bank is selected based on variation in encoder embeddings to represent diverse tissue morphologies. Their selected patch identities are disjoint. Each bank forms an ordered sequence, and corresponding positions determine source-to-reference assignment. The main evaluations assign 2% of eligible patches to each bank. Coverage analyses are reported in Supplementary Methods S15.

$$\mathcal{E} = \{u \in \mathcal{X}_{\text{train}}: \rho(u) \geq 0.50\}, N = |\mathcal{E}| \quad \text{(S1)}$$

$$n = \lfloor pN \rfloor \quad \text{(S2)}$$

$$\mathcal{S} = (t_1, \cdots, t_n), \mathcal{F} = (x_1, \cdots, x_n) \quad \text{(S3)}$$

$$\{t_i\}_{i=1}^{n} \cap \{x_i\}_{i=1}^{n} = \emptyset, t_i, x_i \in \mathcal{E} \quad \text{(S4)}$$

$$\mathcal{P} = \{(x_i, t_i)\}_{i=1}^{n} \quad \text{(S5)}$$

For Style-bank selection, each patch is represented by the mean and standard deviation of the three CIELAB channels, yielding six color measurements. For Fitting-bank selection, each patch is represented by the first 32 principal-component coordinates of encoder embeddings from the corresponding input branch. PCA is estimated independently for each encoder and branch without whitening. Each coordinate is centered and divided by its standard deviation across the eligible population. A zero or numerically negligible standard deviation is replaced by one. These standardized coordinates define distances for bank selection. Displacement estimation and projection are performed in the native encoder embedding space. The number of strata in each standardized space is

$$K = \min(256, N) \quad \text{(S6)}$$

For the reported cohorts $N$ exceeded 256, giving 256 strata. These groups partition the color or embedding coordinate space without biological or source labels. Their number controls sampling priority. Bank size, pair count and response rank are determined separately. Centroids are initialized from deterministically spaced rows in stable order with seed 42. Five Lloyd iterations [3] process the eligible population in the primary analyses. Empty strata are repaired with patches farthest from the populated centroids.

The traversal begins with the largest stratum, then selects the centroid farthest from those already ordered. Stratum size provides a stable tie-break for equal distances. Patches within each stratum are ordered by distance to their centroid and then by stable patch identifier. A round-robin pass takes one candidate from each stratum in that order before advancing to its next candidate. This produces complete Style and Fitting priorities that cover distinct regions of their respective color and embedding spaces before repeatedly sampling any one region.

Joint allocation prevents role overlap. Odd allocation steps give precedence to the Style priority, and even steps give precedence to the Fitting priority. A patch already assigned to one bank is skipped in the other priority. Stable patch identifiers resolve all remaining ties. The retained Style and Fitting orders are paired by position. Increasing p extends the existing banks rather than replacing their members.

**Algorithm 1. Deterministic bank construction and pairing**

| Stage | Operation |
|---|---|
| Input | Eligible patches, complete Style and Fitting priorities, and required bank size n |
| Initialization | Create empty ordered Style and Fitting banks and an empty assigned-patch set |
| Odd allocation step | Append the first unassigned patch in the Style priority, followed by the first unassigned patch in the Fitting priority |
| Even allocation step | Append the first unassigned patch in the Fitting priority, followed by the first unassigned patch in the Style priority |
| Assignment update | Add each appended patch to the assigned-patch set immediately |
| Pairing | Pair patches at the same retained position in the Fitting and Style banks |
| Output | Return disjoint ordered banks and the one-to-one ordered patch pairs |

The allocator was extended to one half of the eligible population to define complete role-specific master sequences. Smaller bank fractions were obtained as nested prefixes of these sequences. A dedicated bank-coverage analysis selected p=0.02, which was then fixed for the reported PathoROB, patch aggregation, TCGA-BRCA, and HUMC evaluations. The disjoint roles therefore expose approximately $2pN = 0.04N$ unique eligible patches across the two banks. Supplementary Data 1 records the estimation scope for each analysis.

#### S4. Paired counterfactual stain-appearance intervention

The intervention defines a controlled alternative stain appearance for one digital tissue field under fixed spatial correspondence. Counterfactual denotes this spatially paired change in digital appearance. Physical staining and scanning are evaluated separately. For pair $i$, the deterministic transformation $\mathcal{T}$ applies the stain appearance of Style patch $t_i$ to Fitting patch $x_i$, yielding the paired stain-appearance variant $\tilde{x}_i$. The fixed encoder then produces the source and counterfactual native embeddings.

$$\tilde{x}_i = \mathcal{T}(x_i; t_i) \quad \text{(S7)}$$

$$z_i = f(x_i), \tilde{z}_i = f(\tilde{x}_i) \quad \text{(S8)}$$

All encoders and both input branches use the same ambient Euclidean displacement and centered orthogonal projection.

$$\delta_i = \tilde{z}_i - z_i \tag{S9}$$

The tilde marks the paired stain-appearance variant or its embedding. The reported experiments instantiated $\mathcal{T}$ with Reinhard stain normalization [4]. Another deterministic stain-appearance transformation can be substituted without changing the subsequent displacement analysis. The Style patch is a patch-level appearance reference. The operator does not require a slide identifier, institution label, or one-reference-per-slide rule.

#### S5. Equal pair contribution

Each retained pair contributes one embedding difference to the displacement matrix with equal weight. Equation (S10) records these weights, and Eq. (S11) gives their corresponding row multipliers.

$$w_i = \frac{1}{n}, i = 1, \dots, n \tag{S10}$$

The resulting general row multiplier is identically one, as shown in Eq. (S11).

$$\sqrt{n w_i} = 1 \tag{S11}$$

Each embedding, displacement and center is defined as a d by 1 column vector. The displacement matrix has n rows and d columns, with one transposed displacement vector per row. All retained pairs enter the matrix with equal coefficients.

$$D = \begin{bmatrix} \delta_1{}^{\mathrm{T}} \\ \vdots \\ \delta_n{}^{\mathrm{T}} \end{bmatrix} \in \mathbb{R}^{n \times d} \tag{S12}$$

With a unit row multiplier for every pair, the unscaled response scatter matrix is

$$D^{\mathrm{T}} D = \sum_{i=1}^{n} \delta_i \, \delta_i{}^{\mathrm{T}} \in \mathbb{R}^{d \times d} \tag{S13}$$

Equations (S10-S13) make the equal-contribution rule explicit while retaining $w_i$ only as bookkeeping notation. The resulting uncentered displacement matrix is supplied directly to singular value decomposition.

#### S6. Response-rank selection

The response spectrum quantifies embedding changes across paired tissue patches. Rank selection determines the leading components included in the projection by comparing their squared singular values with a permutation reference. This reference preserves the distribution within each embedding coordinate while disrupting correspondence across coordinates. Randomized thin singular value decomposition [5] approximates the leading spectrum of D. Let L denote the number of leading components evaluated at the current spectral width.

$$D \approx U_L \Sigma_L V_L{}^{\mathrm{T}} \tag{S14}$$

$$U_L \in \mathbb{R}^{n \times L}, \Sigma_L \in \mathbb{R}^{L \times L}, V_L \in \mathbb{R}^{d \times L}, L \leq \min(n, d) \tag{S15}$$

$$\Sigma_L = \mathrm{diag}(\sigma_1, \cdots, \sigma_L), V_L = [v_1 \cdots v_L] \tag{S16}$$

The diagonal entries of $\Sigma_L$ contain singular values in descending order. Their squares quantify the magnitude of embedding change explained by each component. The left singular vectors describe the corresponding displacement patterns across patch pairs. The columns of $V_L$ are the corresponding orthonormal directions in the $d$-dimensional embedding space.

One hundred permuted displacement matrices are generated. For every replicate $b$ and feature coordinate $j$, $\pi_j^{(b)}$ is sampled independently and uniformly from all permutations of $\{1, \cdots, n\}$. Each permutation shuffles the pair indices within one embedding coordinate. Independent permutations across coordinates generate the reference matrices in Eq. (S17).

$$D_{0,ij}^{(b)} = D_{\pi_j^{(b)}(i),j}, i = 1, \cdots, n, j = 1, \cdots, d, b = 1, \cdots, 100 \quad \text{(S17)}$$

In Eq. (S18), the operator $\text{sv}_L(A)$ returns the leading $L$ singular values of matrix $A$ in non-increasing order. Rank selection requires only these ordered reference singular values.

$$\left(\sigma_{0,1}^{(b)}, \cdots, \sigma_{0,L}^{(b)}\right) = \text{sv}_L\left({D_0}^{(b)}\right), b = 1, \cdots, 100 \quad \text{(S18)}$$

Equation (S19) defines the squared singular values for the observed and permuted displacement matrices. $Q_{0.95}$ is the empirical 95th-percentile operator applied separately to each component across the 100 permuted matrices.

$$\lambda_k = {\sigma_k}^2, \lambda_{0,k}^{(b)} = {\sigma_{0,k}^{(b)}}^2 \quad \text{(S19)}$$

$$q_k = Q_{0.95}\left(\lambda_{0,k}^{(1)}, \cdots, \lambda_{0,k}^{(100)}\right) \quad \text{(S20)}$$

The threshold provides an empirical component-wise reference. Exceedance alone carries no family-wise error guarantee. Equation (S21) defines the positive excess of each observed squared singular value above its reference.

$$e_k = \max(\lambda_k - q_k, 0) \quad \text{(S21)}$$

Rank selection considers consecutive leading components whose squared singular values exceed their references. This interval ends immediately before the first component at or below its reference. The additional value L+1 in Eq. (S22) assigns an endpoint of L if all evaluated components exceed their references.

$$r_s = \min(\{k \in \{1, \cdots, L\}: \lambda_k \le q_k\} \cup \{L + 1\}) - 1 \quad \text{(S22)}$$

For cumulative rank selection, the excess is retained within this leading interval and set to zero outside it.

$$\tilde{e}_k = \begin{cases} \max \quad (\lambda_k - q_k, 0) & \text{if} \quad k \le r_s \\ 0 & \text{if} \quad k > r_s \end{cases} \quad \text{(S23)}$$

DELTA selects the smallest rank accounting for $\eta$ = 0.90 of the summed excess within the leading interval. The selected rank lies between zero and $r_s$.

$$r = \begin{cases} 0 & \text{if} \quad r_s = 0 \\ \min\left\{j \in \{1, \cdots, r_s\}: \frac{\sum_{k=1}^{j} \tilde{e}_k}{\sum_{k=1}^{r_s} \tilde{e}_k} \ge \eta\right\} & \text{if} \quad r_s \ge 1 \end{cases}, \eta = 0.90 \quad \text{(S24)}$$

The spectral search initially evaluates up to 64 components. If every evaluated component exceeds its reference, the search width doubles, up to the smaller of the pair count and embedding dimension. Expansion continues until a component falls at or below its reference or the full available spectrum has been evaluated. If $r_s$ = 0, then $r$ = 0 and the transformation is the identity. Zero-rank estimates are recorded separately from successful subspace estimates. The TCGA-BRCA survival workflow treats these estimates as fitting failures, as specified in S12. Each input branch has an independently estimated response basis and selected rank.

Subspace stability is assessed by re-estimating the response basis across 200 block-bootstrap replicates. The unsupervised Fitting strata defined in Supplementary Methods S3 form the resampling units, and the selected rank remains fixed throughout this assessment. The bootstrap index $s$ ranges from 1 to 200, and $b$ indexes the permutation replicates. For a positive selected rank, the starred basis in Eq. (S25) is the $d$ by $r$ basis estimated from bootstrap replicate $s$, and $B$ is the basis estimated from the complete displacement matrix. The Frobenius norm is the square root of the sum of squared matrix entries. The normalized projection overlap is

$$\mathrm{Overlap}\left(B^{*(s)}, B\right) = \frac{\left\|B^{*(s)^{\mathrm{T}}}B\right\|_F^2}{r} \tag{S25}$$

Normalized projection overlap equals one for identical subspaces and zero for orthogonal subspaces. Rank-zero estimates receive a not-applicable stability value. Overlap values are recorded as diagnostics of subspace stability. The original banks, selected rank and operator are retained for downstream evaluation. Stochastic numerical procedures for DELTA operator estimation are initialized with seed 42.

#### S7. DELTA projection

The selected response basis contains the first $r$ right singular vectors and has $d$ rows and $r$ orthonormal columns. In Eq. (S26), $I_r$ is the $r$ by $r$ identity matrix. The stored center is the unweighted arithmetic mean of the $n$ source Fitting representations.

$$B = [v_1 \cdots v_r] \in \mathbb{R}^{d\times r}, B^{\mathrm{T}}B = I_r \tag{S26}$$

$$\mu = \frac{\sum_{i=1}^{n} z_i}{n} \in \mathbb{R}^{d\times 1} \tag{S27}$$

Every pair contributes equally to the response spectrum, and the center is the arithmetic mean of the same retained source Fitting representations. For a new $d$ by 1 column-vector embedding $z$, $I_d$ is the $d$ by $d$ identity matrix and $BB^{\mathrm{T}}$ is the orthogonal projector onto the selected response subspace. DELTA applies

$$z_{\Delta} = \mu + (I_d - BB^{\mathrm{T}})(z - \mu) = z - BB^{\mathrm{T}}(z - \mu) \tag{S28}$$

Equation (S28) defines a centered orthogonal projection in the Euclidean embedding space. The operator stores $\mu$, $B$, and the applied rank. The stored operator is applied to new embeddings after the preprocessing prescribed for its input branch. The center and basis remain fixed during inference. The projection preserves the mean of the original Fitting bank embeddings and the embedding dimension.

#### S8. Comparison methods

The raw encoder embedding was the reference representation. PathoROB, patch aggregation and TCGA-BRCA evaluations applied Reinhard normalization [4] toward an external reference derived from the averaged color statistics of 500 TCGA-LUSC patches from 250 patients outside the evaluation cohorts. HUMC selected a separate reference within each fold. One representative patch from each training non-tumor case contributed CIELAB channel means and standard deviations. Median and median-absolute-deviation standardization defined a robust descriptor space, and the patch closest to its center supplied the reference appearance.

ComBat estimates feature-wise location and scale parameters conditional on an observed batch label [6,7]. ComBat, evaluated under a transductive protocol, accesses the evaluation batch and is included in the patch-level and patch-aggregated comparison families specified in S14. Domain-adversarial training optimizes a task predictor and domain discriminator through gradient reversal [8]. It requires domain labels and task labels during training. DELTA requires neither label type for operator estimation.

#### S9. PathoROB representation analysis

The descriptive representation analysis retained four multicenter collections from PathoROB [2] to assess biological organization and acquisition-source sensitivity. Camelyon [9] combined CAMELYON16 and CAMELYON17 and contained 22,402 normal and tumor patches from 97 lymph node slides across five medical centers. TCGA4×4 [10,11] contained 8,160 patches from 272 slides representing 272 patients across eight medical centers and four cancer classes from breast, colon and lung tissues. Tolkach ESCA [12,13] contained 16,300 patches from 77 esophageal slides covering six tissue classes across three medical centers and an external cohort. TCGA2×2 [10,11] contained 59,250 distinct patches from 1,975 slides spanning 21 cancer classes. Operators were estimated from the eligible collections without predictive train–test partitions. Endpoint-specific populations then followed PathoROB. Camelyon neighborhood analysis excluded out-of-domain (OOD) sources and retained 20,400 patches. Tolkach neighborhood analysis retained 9,000 patches from 43 slides and three sources, with 500 patches in each source–class cell. TCGA2×2 evaluated its 94 prescribed comparison subsets. These evaluation populations are narrower than the released image collections in some datasets.

TCGA4×4 and TCGA2×2 shared 6,720 patch identities from 224 slides. The 112,800 TCGA2×2 comparison memberships include repeated appearances of its 59,250 distinct patches. The aggregation experiments reused TCGA4×4, and the NSCLC task was nested within it. In addition, 68 TCGA4×4 patients and 245 TCGA2×2 patients occurred in the TCGA-BRCA survival cohort. The tasks assess different outcomes and sampling regimes within partly overlapping public material.

Biological nearest-neighbor performance, confounder insensitivity and robustness index were averaged equally across Camelyon, TCGA2×2 and Tolkach ESCA. These neighborhood analyses provided the summaries in Source Data. Clustering score averaged Camelyon, TCGA4×4 and Tolkach ESCA. DANN and DANN after Reinhard normalization neighborhood summaries averaged Camelyon and Tolkach ESCA. Their clustering summaries retained the three-dataset scope. For Fig. 2, representation metrics were averaged equally across Camelyon, TCGA4×4 and Tolkach ESCA for each encoder and method. Dataset-specific neighborhood analyses retained TCGA2×2.

Neighbor search operated on L2-normalized embeddings and excluded every candidate patch from the query slide. Euclidean distances in this space give the same neighbor order as cosine similarity. The fixed neighborhood sizes were 11 for Camelyon, 61 for TCGA and 46 for Tolkach ESCA. Biological nearest-neighbor performance was the balanced accuracy of the majority biological label among the retained neighbors. Confounder insensitivity [2] was the pooled number of other-source neighbor incidences divided by the pooled number of same-source incidences, irrespective of biological class. This ratio is not bounded by one. Robustness index [2] was the number of same-biology and different-source incidences divided by the sum of those incidences and different-biology and same-source incidences. Both ratios pooled query–neighbor counts without logarithms, clipping or averaging individual patch ratios.

TCGA2×2 constructed neighborhoods separately in each comparison subset. Biological-label balanced accuracies were averaged across the 94 subsets. Confounder insensitivity and robustness index pooled subset-specific neighbor incidences, retaining repeated membership according to the benchmark structure. K-means clustering operated with cosine distance. Silhouette score [14] selected the number of clusters from 2 to 30, and the final solution had 50 initializations. The clustering score subtracted adjusted Rand agreement [15] with source labels from adjusted Rand agreement with biological labels [2].

Labels defined the evaluation endpoints. DELTA operator estimation was label-free. On embeddings from Reinhard-normalized images, DELTA increased the mean robustness index from 0.7465 to 0.7753 across Camelyon, TCGA2×2 and Tolkach ESCA. The paired gain was 0.0288 with a 95% confidence interval of 0.0200 to 0.0381. Nineteen of 20 encoders improved, and the Holm-adjusted exact sign-flip P value was 0.0001144.

### S10. Patch classification

Patch classification retained the official PathoROB source partitions [2]. Camelyon training and in-domain (ID) evaluation drew on RUMC and UMCU, while CWZ, LPON and RST supplied OOD images. TCGA4×4 and Tolkach ESCA similarly separated development and OOD sources. Twenty repetitions at eight, seven and four source–class association settings yielded 160 Camelyon, 140 TCGA4×4 and 80 Tolkach splits. These settings changed the class composition associated with each training source. The task labels were normal versus tumor for Camelyon, four cancer types for TCGA, and six tissue classes for Tolkach. Held-out class counts were not uniformly equal, including the absence of regression-tissue patches from the TCGA source of Tolkach.

Each representation was evaluated with logistic regression on identical split memberships. Validation balanced accuracy selected the inverse regularization parameter from 15 logarithmically spaced values between $10^{-8}$ and $10^{4}$. Training embeddings alone estimated the selected classifier, and DELTA estimation was confined to the same training partition. Balanced accuracy was the arithmetic mean of recall across classes represented in the true labels of the evaluated partition.

All four partitions were patch-disjoint. Camelyon and TCGA4×4 were also slide-disjoint. The retained Tolkach procedure could assign different 100-patch blocks from one case to training and validation. Its test cases remained slide-disjoint from both development partitions. The separate aggregation experiment confined each available patient-group identifier to a single partition.

Average performance drop [2] summarized relative balanced accuracy under varying center–class associations. For each encoder, dataset and evaluation domain, the reference was RAW balanced accuracy at the balanced condition averaged across repetitions. At each subsequent condition, the difference between method-specific balanced accuracy and this fixed reference was divided by the reference. These relative changes were averaged across repetitions and subsequent conditions. Negative values indicated accuracy below the RAW balanced-condition reference. Results were averaged over 20 repetitions within dataset and equally across the three datasets within each encoder. The 20 encoders formed the paired statistical units. After DELTA, mean ID average performance drop changed from −0.0529 to −0.0380. The paired gain was 0.0149 with a 95% confidence interval of 0.0110 to 0.0192 and a Holm-adjusted exact sign-flip P value of $2.29 \times 10^{-5}$. All 20 encoders improved. The OOD mean changed from −0.0345 to −0.0212. Its paired gain was 0.0133 with a 95% confidence interval of 0.0088 to 0.0183 and a Holm-adjusted P value of $5.15 \times 10^{-5}$. Nineteen encoders improved.

#### S11. Patch aggregation

The aggregation experiment assessed slide- and patient-level classification from the Camelyon [9] and TCGA4×4 [10,11] patch collections, retaining all released patches. Camelyon comprised 49 normal and 48 tumor slide-level patch sets. Its 91 split-group identifiers consisted of 68 CAMELYON16 slide-based proxies and 23 CAMELYON17 patient barcodes covering 29 slides. All patch sets with the same available identifier remained in one partition. TCGA patient identifiers were defined by the first 12 slide-barcode characters, with one slide per patient in these collections. Each of the four cancer classes contained 68 patients. The NSCLC task retained the same 68 lung adenocarcinoma and 68 lung squamous cell carcinoma patients from the four-class cohort. The binary NSCLC task assessed histological discrimination within lung tissue, complementing the four-class task spanning breast, colon and lung cancers.

Twenty repeated partitions at source–class association levels of 0, 0.25, 0.50, 0.75 and 1.00 produced 100 splits per task. One group from each ID source–class cell was reserved for validation. Training groups followed the prescribed association, remaining ID groups formed the ID test set, and OOD sources formed the OOD test set. Camelyon contained 28 training, 4 validation, 36 ID test and 23 OOD test groups. Corresponding patient counts were 40, 8, 48 and 40 for NSCLC, and 32, 16, 144 and 80 for four-class TCGA.

Slide- or patient-level patch sets formed the bags for multiple-instance learning. A gated-attention multiple-instance model [16] mapped each patch embedding to 256 features through a linear layer, ReLU activation and dropout of 0.25. Parallel tanh and sigmoid branches generated attention scores, which were normalized by softmax. A linear classifier received the weighted bag representation. Training minimized cross-entropy with stochastic gradient descent at learning rate 0.002 for 200 epochs and batches of 32 bags. Validation balanced accuracy selected the epoch checkpoint and weight decay from 0 and 0.01. Architecture, optimization settings and the procedure for validation-based model selection were held constant across representation branches. DELTA operators were estimated exclusively from training patches and applied before attention aggregation. For DANN comparisons, the acquisition-source discriminator received the 256-dimensional pooled bag representation. It comprised a linear layer with 128 outputs, ReLU activation and a linear output layer for the training centers. Biological and acquisition-source cross-entropy losses were summed. The gradient-reversal coefficient increased linearly from zero to one across the 200 training epochs. Optimization and validation-based model selection followed the MIL procedure described above.

Balanced accuracy was the primary endpoint. Macro F1, AUROC, AUPRC, Brier score [17] and expected calibration error [18] with 10 bins were secondary endpoints. Binary AUROC and AUPRC were calculated from positive-class probabilities, with AUPRC calculated as average precision. Four-class AUROC averaged class-specific one-versus-rest values equally, and AUPRC averaged class-specific average precision equally. Binary Brier score was the mean squared error of positive-class probabilities. Four-class Brier score summed squared probability errors across classes before averaging across slide- or patient-level patch sets. Expected calibration error grouped maximum predicted class probabilities into 10 equal-width intervals over zero to one. Each interval contributed its sample fraction multiplied by the absolute difference between mean confidence and classification accuracy. Results were averaged over the 100 splits within each task and encoder (Fig. 3 and Supplementary Fig. 2). The three task means received equal weights before paired inference across encoders. Confidence intervals, statistical tests and multiplicity adjustment are detailed in Supplementary Methods S14.

#### S12. TCGA-BRCA WSI-level survival analysis

The TCGA-BRCA cohort [19] contained 996 patients and 1,057 slides, yielding 6,969,941 tissue tiles. The initial cohort contained 1,009 patients, of whom 13 lacked entries in the frozen WSI analysis index. GDC vital status and days to death [20] defined observed deaths. For patients without a recorded death, the maximum positive follow-up time defined censoring. Patient identifiers were the first 12 TCGA barcode characters, and center denoted the tissue-source-site code. All slides from one patient remained in a single partition.

Six tissue-source sites contributed training, validation and ID test patients. Their TCGA codes were A2, A8, AR, BH, D8 and E2. Patient assignments followed a frozen split generated with seed 7028 and remained fixed across encoders and optimization seeds. Fifteen other sites formed the OOD test. The four partitions contained 381, 84, 80 and 451 patients, with 61, 14, 11 and 55 deaths, respectively. DELTA corrected tile embeddings before averaging all eligible tiles from all slides belonging to a patient. Feature standardization was estimated from training patients.

The eligible training population contains 2,799,421 tiles. Each bank contains 55,988 tiles at p=0.02, producing 55,988 paired interventions for each encoder and input branch. Every retained pair enters rank estimation once and with the same coefficient, under the rule used in all DELTA analyses. The adaptive singular-spectrum analysis starts at width 64 and has no fixed row or rank cap. Outcome, event time, center, validation embeddings, and test embeddings do not enter operator estimation. In TCGA-BRCA survival analysis, zero-rank operator estimates were treated as fitting failures and did not generate a reportable DELTA comparison.

The survival learner is a neural Cox risk model [21] with a 256-unit hidden layer, ReLU activation, dropout of 0.20, and a scalar risk output. AdamW optimization [22] uses a learning rate of 0.001 and weight decay of 0.0001. The model minimizes the Efron partial-likelihood loss [23] for up to 200 epochs. Each epoch comprises one full-batch update over all 381 training patients, with risk sets defined across the complete training cohort. Early stopping monitors validation Uno C-index with a patience of 20 epochs. Models are fitted with seeds 7028, 17028, and 27028, and patient risk predictions are averaged across seeds before model-level inference. Seeds quantify optimization variability and are not statistical units.

Uno C-index [24] is the primary endpoint. The censoring Kaplan-Meier distribution [25] is estimated from training patients, and the truncation time is the maximum training event time. Time-dependent AUC [26] is evaluated at the 25th, 50th, and 75th percentiles of training event times. Mean time-dependent AUC is the arithmetic mean of the three time-specific AUC values. Integrated Brier score [27] uses the unique values from 20 quantiles between 0.10 and 0.90 of the training event-time distribution and trapezoidal integration normalized by the evaluation interval.

The Breslow baseline hazard [28] is estimated once from the training patients using seed-averaged training risk and is not re-estimated in either test domain. Harrell C-index [29], integrated Brier score, and mean time-dependent AUC are secondary endpoints (Fig. 4a,c and Supplementary Fig. 3a,c). Negative DELTA-minus-comparator differences in integrated Brier score indicate a reduction in error. The same training-derived risk reference is used for baseline-hazard estimation and test survival probabilities and is never recomputed from a test cohort or batch. For integrated Brier score, event contributions are weighted by the inverse censoring probability immediately preceding the event. Contributions from patients surviving beyond an evaluation time are weighted by the inverse censoring probability at that time.

#### S13. HUMC normative screening

The HUMC cohort comprised gastric biopsy specimens collected between December 2024 and December 2025 from 132 distinct patients, including 100 non-tumor cases and 32 biopsy-proven cancer cases. Training non-tumor images in each fold defined a reference distribution of tissue embeddings against which held-out cases were scored. The DELTA operator, normative reference and threshold were estimated within that fold. Cancer images and labels were excluded from all three steps. The primary analysis applied five Lloyd iterations to each bank, with every iteration processing the full eligible training pool. The primary result tables retain the five-iteration analysis.

The frozen inventory retained patches with tissue fraction of at least 0.50. Each case had at least 50 eligible patches and contributed at most 2,000 sampled patches. Cases below the upper limit retained all eligible patches. Larger cases received proportional slide quotas followed by deterministic, evenly spaced sampling from coordinate-ordered patches. The resulting 117,277 patches were shared across representation branches. Case identifiers were assigned to five folds by deterministic hashing with seed 42, separately within non-tumor and cancer groups. Each fold held out 20 non-tumor cases and six or seven cancer cases, leaving 80 non-tumor cases for estimation. Multiple slides from one case remained in the same fold.

A patch anomaly score was the mean cosine distance to its 20 nearest training non-tumor embeddings. The case score averaged the highest 1% of patch scores, with at least five patches. This upper-tail aggregation targets focal tissue departures that could be diluted by averaging an entire biopsy. Leave-one-case-out scoring excluded every patch from the query case from the nearest-neighbor reference. The fold-specific DELTA operator and Reinhard reference remained fixed during this step. Each representation branch set its threshold at the linearly interpolated 95th percentile of its own leave-one-case-out training non-tumor case scores. The threshold remained fixed for test non-tumor and cancer cases. Scores at or above the threshold were classified as positive. Specificity at complete cancer sensitivity and sensitivity at complete non-tumor specificity were exploratory extreme operating points.

Endpoints were calculated separately for each encoder from the pooled out-of-fold scores of all 132 cases. AUROC summarized ranking across case scores, and AUPRC was calculated as average precision under the observed cancer prevalence [30]. Sensitivity and specificity were calculated from pooled case predictions at the corresponding fold-specific thresholds. Encoder-level endpoints were then averaged across the 20 FMs (Fig. 4b,d and Supplementary Fig. 3b,d). The training non-tumor pool supplied images for label-free operator estimation and cases for normative-reference construction and threshold selection. All case scores and thresholds were fixed before cancer status entered endpoint calculation.

### S14. Statistical analysis

The foundation model is the paired statistical unit for patch-level, patch-aggregated and WSI-level survival comparisons. Dataset or task results are averaged within encoder before inference. Panel intervals describe variation in paired effects across the evaluated models on the fixed cohorts, not patient-sampling uncertainty or independent replication by 20 patient cohorts. Shared model ancestry can also limit encoder independence. HUMC additionally incorporates paired case resampling as described below. PLISM inference retains tissue groups as the paired units with the encoder panel fixed, as detailed in S16.

Patch representation and patch-classification confidence intervals were calculated from 10,000 bootstrap resamples [31] of the 20 paired model effects. Dataset-level results were averaged within each encoder before inference. Two-sided exact sign-flip tests evaluated comparisons with reference methods, including ComBat. Holm adjustment [32] was applied across seven comparators for each representation metric and DELTA input branch in Fig. 2. Comparators included RAW, DANN and ComBat in each input branch and DELTA in the other input branch. Dataset-specific analyses in Fig. 2 were adjusted across 24 contrasts comprising eight endpoints and three datasets within each DELTA input branch. Each classification family comprised 12 contrasts across the ID and OOD domains and six comparators for one metric. These were raw embeddings, DANN and ComBat in each of the original and Reinhard input branches. The separate representation summaries described in S9 retained Holm adjustment across 16 contrasts comprising four endpoints and four comparators. These were raw encoder embeddings, Reinhard normalization, ComBat applied to original embeddings and ComBat applied to Reinhard-normalized embeddings. DANN summaries for these analyses remained descriptive. Wilcoxon signed-rank tests [33] were retained as corroborative analyses.

For patch aggregation, results were averaged across 100 matched splits within each task and then equally across the three tasks within each encoder. DELTA in the original input branch was compared with raw encoder embeddings, Reinhard normalization alone, DANN and ComBat. DELTA after Reinhard normalization was compared with raw encoder embeddings, Reinhard normalization alone, DANN after Reinhard normalization and ComBat after Reinhard normalization. ComBat was evaluated under a transductive protocol in both input branches. Pointwise 95% percentile confidence intervals were calculated from 10,000 bootstrap resamples of the 20 paired encoder-level differences. Two-sided exact Wilcoxon signed-rank tests evaluated each contrast after rounding differences to 12 decimal places, assigning average ranks to ties and excluding zero differences. Estimates and confidence intervals were calculated from unrounded differences. Holm adjustment covered four comparators separately for each DELTA input branch, endpoint and ID or OOD domain. Analyses of individual tasks were adjusted across the four comparators and three tasks. Bootstrap confidence intervals for results averaged across tasks were generated with seed 42. For individual tasks, seeds were determined by task, comparator, input branch and evaluation domain. Source Data reports the direct DELTA–ComBat comparisons.

For WSI-level survival analysis, endpoints were calculated from patient risk predictions averaged across optimization seeds within each encoder. Pointwise 95% percentile confidence intervals were estimated from 20,000 bootstrap resamples of paired encoder-level differences, with the patient cohort held fixed. Two-sided exact sign-flip tests evaluated prespecified contrasts. Holm adjustment covered four comparators within each endpoint, evaluation domain and DELTA input branch. DELTA was compared with raw encoder embeddings, Reinhard normalization, DANN and ComBat. DELTA after Reinhard normalization was compared with raw encoder embeddings, Reinhard normalization alone, DANN after Reinhard normalization and ComBat after Reinhard normalization. Common-control maxT inference [34] provided a multiplicity sensitivity analysis. For this analysis, DELTA was compared with raw encoder embeddings, DANN, Reinhard normalization and DANN after Reinhard normalization. The family for DELTA after Reinhard normalization additionally included DELTA. Each encoder received a common sign across all contrasts within a family. The largest absolute studentized mean difference defined the family-wise reference distribution.

Primary HUMC screening inference comprised 20,000 hierarchical paired-bootstrap replicates. Non-tumor and cancer cases were resampled separately with shared case weights across all methods and encoders, followed by resampling of the 20-encoder panel. Fold-specific thresholds and predictions remained fixed. Two-sided P values were obtained from the centered hierarchical difference distribution. Holm correction covered five comparisons within each metric. The contrasts compared DELTA with raw encoder embeddings and Reinhard normalization, and DELTA after Reinhard normalization with raw encoder embeddings, Reinhard normalization and DELTA. Model-level sign-flip tests were retained as secondary analyses. This inferential resampling was separate from the label-free Fitting-stratum bootstrap that only assessed subspace stability. Missing results were not imputed.

#### S15. Coverage sensitivity analysis

The main analyses fix p = 0.02, a practical operating point chosen during bank-coverage development. The coverage grid contains p = 0.01, 0.02, 0.05, 0.10, 0.20, 0.30, 0.40 and 0.50. Deterministic priorities make larger banks extend preceding prefixes, subject to the disjoint joint-allocation rule. This reduces changes in patch membership between coverage levels. The p = 0.02 setting is not asserted to maximize every endpoint or every backbone. All eight fractions were evaluated. TCGA-BRCA included CONCH, MUSK, H0-mini, Original UNI and Virchow2 in both input branches, giving 80 encoder–branch–fraction conditions. PathoROB included 20 encoders, both branches and Camelyon, TCGA4×4 and Tolkach ESCA. Representation operators were estimated from eligible benchmark cohorts, and classification operators from each split's training patches (Supplementary Fig. 1).

Coverage is defined relative to the eligible patch pool for each analysis. At p = 0.02, the two disjoint banks jointly contain approximately four percent of that pool and one-to-one pairing yields approximately two percent as many paired interventions as patches in that pool. These percentages quantify participation in paired interventions. Total computation also includes encoder extraction and descriptor construction. Coverage evidence is interpreted as development and sensitivity analysis. It does not provide an independent confirmatory evaluation of the selected operating point. Larger banks did not consistently improve downstream performance. In the five-encoder TCGA analysis, mean ID Uno C-index in the original-input DELTA branch was 0.6200, 0.6139 and 0.6064 at fractions of 0.01, 0.02 and 0.50. Corresponding Reinhard-branch means were 0.5807, 0.5783 and 0.5932. The operating fraction was fixed at 0.02 in HUMC without a separate HUMC fraction sweep. At a fixed fraction, bank membership, response-basis estimation and rank selection did not access outcomes.

#### S16. Response-basis controls and PLISM acquisition transfer

PLISM [35] provided registered images of formalin-fixed, paraffin-embedded tissue microarrays representing 46 human tissue categories. The analysis included 13 staining conditions and seven scanners, excluding smartphone images (Supplementary Fig. 4). Within each staining condition, scanner comparisons involved repeated images of an identical physical section. Comparisons across staining conditions involved registered regions on serial sections and therefore included local tissue differences. The H&E condition with Mayer hematoxylin, designated MY in PLISM, served as the fixed staining reference for the scanner analysis. This subset provided 3,417 corresponding tissue regions imaged by all seven scanners. A tissue region was the retrieval unit.

DELTA and comparison operators were estimated from TCGA-BRCA for original-input embeddings from 20 FMs and applied to PLISM without refitting. Operator estimation and rank selection excluded PLISM. For each encoder, the alternative operators retained the projection center and selected rank of the DELTA operator and applied centered orthogonal projection. The Fitting-bank PCA basis comprised the leading principal components of centered original Fitting-bank embeddings. The unpaired displacement basis followed a fixed derangement of the original partners of transformed embeddings, with no original pair retained. Five random orthonormal bases were generated from Gaussian random vectors with seed 42 through QR decomposition. Distance and retrieval endpoints were calculated separately for each random orthonormal basis and then averaged across the five projections.

Distance analyses included 273 acquisition-condition pairs differing only in scanner, 546 differing only in staining condition and 3,276 differing in both. Squared embedding-distance reduction followed Eq. (10). Ratios were calculated for raw squared embedding distances greater than $10^{-12}$. In the reported summaries across all acquisition conditions, matched-region pairs with raw squared distances at or below this threshold contributed zero reduction. Such pairs were omitted from the MY-only distance summaries. Raw encoder embeddings corresponded to the identity operator and had zero reduction.

Retrieval compared patches from each evaluated acquisition condition with candidate patches from a designated reference condition. The NanoZoomer-S60 scanner (S60) and MY served as the reference scanner and staining condition, respectively. Retrieval comprised 78 scanner comparisons against S60 within each staining condition, 84 staining comparisons against MY within each scanner, and 72 combined comparisons against patches acquired under MY staining with S60.

Matched-field top-1 retrieval measured the proportion of evaluated patches for which the corresponding reference patch ranked first. Within-tissue top-1 retrieval restricted candidates to the tissue category of the evaluated patch. In the MY scanner analysis, S60 patches were ranked by Euclidean embedding distance for each patch from the other six scanners. Reciprocal rank was the inverse rank of the corresponding S60 patch. Tissue-category concordance excluded the corresponding patch and measured agreement in tissue category with the nearest remaining patch (Supplementary Fig. 5). These endpoints distinguished recovery of the corresponding region from preservation of tissue-category neighborhoods.

Within the MY subset, distance reductions were averaged across corresponding regions and 21 scanner pairs within each tissue category. Retrieval endpoints were averaged across regions and the six scanners evaluated against S60 within each category. Category means were then averaged within each encoder and across the 20 FMs. These results were summarized descriptively in Supplementary Table 4.

Across all acquisition conditions, region-level endpoints were averaged within each tissue group and then equally across the 46 groups for each acquisition comparison. Scanner contrasts were averaged across comparisons within each staining condition and then equally across the 13 conditions. Staining contrasts were averaged within each scanner and then equally across the seven scanners. Contrasts differing in both factors averaged all relevant comparisons equally. This procedure was applied separately to distance and retrieval endpoints with their respective comparison sets. Final summaries equally weighted the 20 FMs. Paired differences between methods were averaged across the 20 FMs within each tissue group. Pointwise 95% percentile confidence intervals were calculated from 20,000 paired bootstrap resamples of the 46 tissue groups, with the FMs held fixed. Resampling preserved pairing across methods, endpoints and acquisition comparisons, with seed 42. Two-sided conditional exact Wilcoxon signed-rank tests assigned average ranks to ties and excluded zero differences. Paired differences were rounded to 12 decimal places for hypothesis testing before conversion to percentage points for display. Estimates and confidence intervals were calculated from unrounded differences. Holm adjustment covered 36 comparisons across three endpoints, three acquisition contrasts and four comparators. Bonferroni adjustment across these 36 comparisons provided a sensitivity analysis. Inference was conditional on the observed tissue groups, acquisition conditions and FMs. Bootstrap resampling treated tissue groups as exchangeable. Signed-rank tests assumed independent differences with a symmetric distribution under the null hypothesis. Results across acquisition conditions are provided in Supplementary Data 1 and Source Data.

Scanner and staining-condition prediction was evaluated with linear ridge-classification probes including intercepts. Five cross-validation folds partitioned the 46 tissue groups, with all regions and acquisition conditions from each group assigned to one fold. Each training group received equal total weight, with acquisition classes balanced within group. The ridge coefficient was set to 0.01 times the mean diagonal of the RAW training covariance and applied to all methods within each fold. Balanced accuracy was calculated within each held-out tissue group and then averaged across tissue groups and the 20 FMs. Results from the five random orthonormal bases were averaged before comparison. Pointwise 95% percentile confidence intervals were calculated from 20,000 paired bootstrap resamples of tissue groups, with the fitted classifiers and their out-of-fold predictions held fixed. Two-sided conditional exact Wilcoxon signed-rank tests followed the procedure above, with Holm adjustment across eight comparisons comprising two prediction targets and four comparators. Lower balanced accuracy indicated lower accuracy in predicting scanner identity or staining condition.

### S17. STP-BENCH spatial molecular prediction

Spatial gene-expression prediction evaluated the retention of molecular information after DELTA correction. The analysis followed the official STP-BENCH linear-probe protocol implemented in LinearProb with the frozen UNI-v2 encoder [1]. Embeddings predicted measured expression at corresponding tissue regions. Visium supplied capture spots, and Xenium single-cell measurements were pooled upstream into comparable pseudo-spots. Visium and Xenium measurements covered breast, prostate, renal, lung, brain and pancreatic cancers. The seven internal datasets contributed 202 sections across 33 fixed folds. The two datasets from HEST-1k [36], HEST-PRAD and HEST-CCRCC, contributed two and six folds, respectively. WUSTL-BRCA [37], NCCHE-Xenium and NCCHE-Visium [38], SNU-GBM [39] and WUSTL-PDAC [37] each had five folds. Supplementary Data 1 identifies the cancer type, platform and section count of each dataset. Counts represent sections rather than independent patients.

The six external datasets specified in STP-BENCH contributed 65 sections. The fixed fold-0 source predictor was transferred from WUSTL-BRCA to HEST-BRCA and Massey-TNBC [40], from NCCHE-Xenium to HEST-LUAD, from HEST-CCRCC to WUSTL-RCC, from SNU-GBM to HEST-GBM and from WUSTL-PDAC to HEST-PAAD. Prediction targets comprised the 200 high-mean highly variable genes (HMHVG) and 16 tumor-microenvironment markers specified by the benchmark. Additional transfers from WUSTL-BRCA evaluated 68 sections from 23 patients in ST-Net [41] and 36 sections from eight patients in HER2ST [42]. For these two cohorts, evaluation included genes in the trained gene panels and present in every section of the target cohort. Gene names were matched by approved symbols from the HUGO Gene Nomenclature Committee (HGNC). DELTA operators and predictors remained fixed during external transfer. This experiment examined UNI-v2 only, separately from the 20-encoder evaluations elsewhere in the study.

The LinearProb implementation was evaluated at repository revision dd476415. The frozen UNI-v2 encoder produced 1,536-dimensional embeddings from 224-pixel image regions. Expression targets were log1p-transformed raw counts without counts-per-million normalization or smoothing. Official training folds, optimization settings and early stopping were retained. Separate linear predictors were trained on raw encoder embeddings and DELTA-corrected embeddings, with representation as the only difference between the two evaluations.

Within each source-training fold, the eligible population comprised image regions retained by STP-BENCH patch quality control without an additional DELTA-specific tissue-fraction threshold. The 2% Style and 2% Fitting banks were disjoint. All paired embedding displacements contributed equally to subspace estimation without a displacement-row cap. Each branch followed rank selection based on 90% of the summed excess above the permutation references, followed by centered orthogonal projection. Expression values, validation data and test data were excluded from bank selection, response-basis estimation, rank selection and projection centering. The source operator remained fixed during internal validation. For external transfer, the source operator and trained linear predictor from fold 0 were applied without target-specific operator estimation, predictor fitting or cohort-statistic calculation.

Pearson correlation quantified linear correspondence between predicted and measured expression, and mean absolute error quantified prediction error on the transformed expression scale. Both metrics were calculated separately for each gene across spots within each section and macro-averaged across evaluable genes (Supplementary Fig. 6). Undefined correlations were omitted through the benchmark torch.nanmean implementation. For internal evaluation, section-level results were averaged within fixed folds and then across folds for descriptive task summaries. Paired changes in Pearson correlation were expressed as DELTA minus raw encoder embeddings. Changes in mean absolute error were expressed as raw encoder embeddings minus DELTA, with positive values indicating improvement for both metrics. Recalculation from stored predictions reproduced official task means to within 1.84 x 10^-7.

Each dataset or external transfer was analyzed separately for each gene panel and metric. Tissue sections within fixed folds constituted the paired sampling units in the original STP-BENCH analyses. Pointwise 95% percentile confidence intervals were calculated from 20,000 paired bootstrap resamples of sections, stratified by fold and averaged with equal fold weights. Two-sided paired section sign-flip tests involved exact enumeration for at most 20 sections and 100,000 Monte Carlo repetitions otherwise. Holm correction was applied across tasks separately for internal validation and the original external transfers within each gene panel and metric. For ST-Net and HER2ST, point estimates averaged section-level metrics with equal section weights. Bootstrap resampling selected patients and retained all sections from each sampled patient. Exact sign-flip tests assigned one shared sign to all sections from each patient. Holm correction was applied across these two additional transfers separately for each gene panel and metric. Patient-level scatter plots averaged section metrics within each patient.

#### S18. Computational environment

Patch-level classifier training and spatial gene-expression model training were performed in the Python 3.9.23 environment. The primary HUMC runs for operator estimation and normative screening were performed in the Python 3.10.18 environment. Supplementary Table 5 summarizes the software configurations documented for these environments.GPU computations were distributed across NVIDIA A100, H100, RTX A6000, RTX 6000 Ada Generation and GeForce RTX 3090 devices under Slurm.

### Figures and Figure Legends

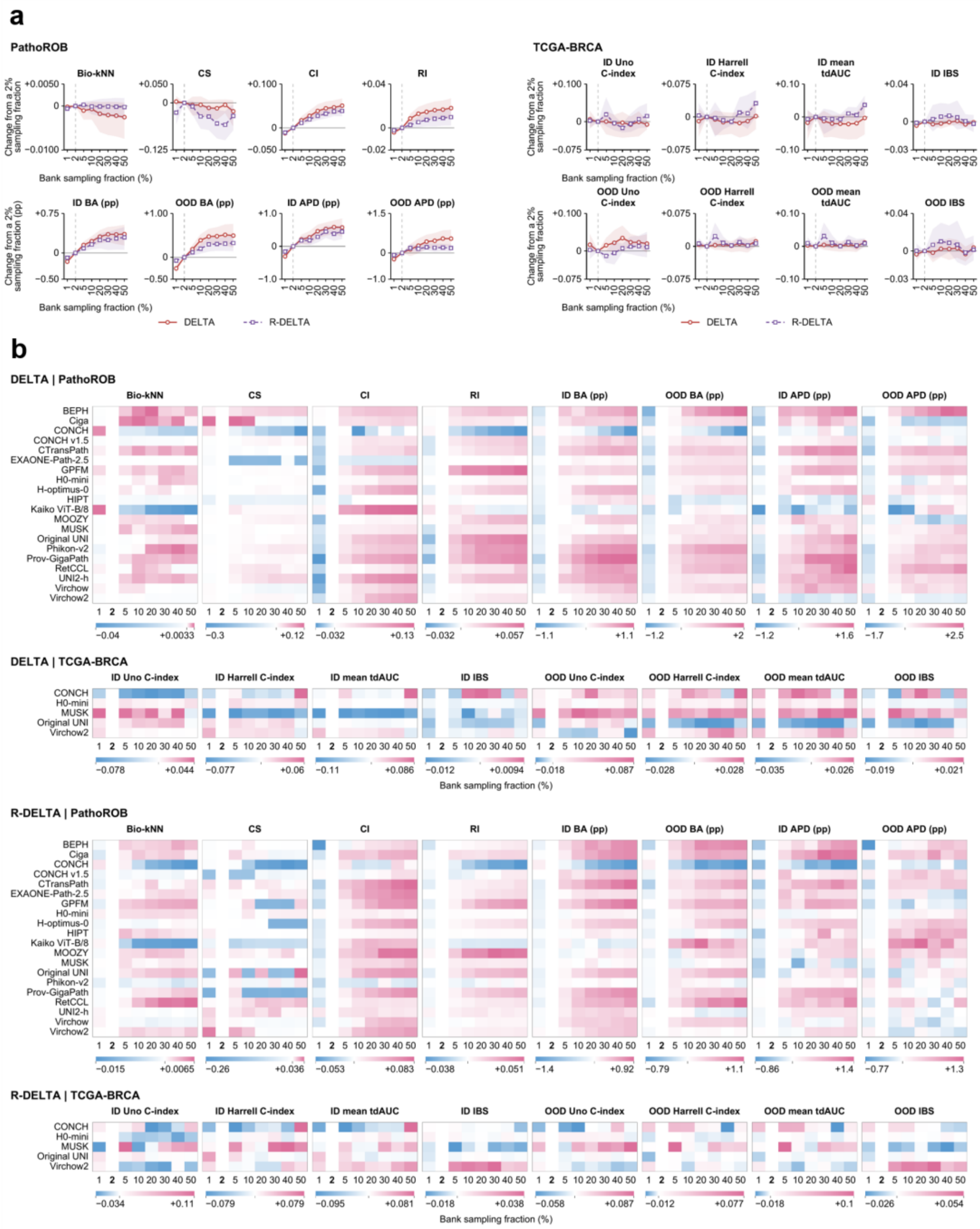


**Supplementary Figure 1. Sensitivity of DELTA to the bank sampling fraction.**

Sampling fraction denotes the proportion of eligible patches selected for each of the disjoint Style and Fitting banks. All values represent changes from the corresponding 2% result within each foundation model (FM) and input branch. **a**, Mean changes across 20 FMs evaluated on PathoROB and five FMs evaluated in TCGA-BRCA survival analysis. PathoROB results were averaged across datasets within each FM. Red solid and purple dashed curves indicate DELTA applied to raw embeddings and Reinhard-normalized embeddings (R-DELTA), respectively. Shaded bands indicate pointwise 95% percentile confidence intervals from 10,000 bootstrap resamples of FMs with replacement, preserving pairing across sampling fractions and input branches. FMs formed the resampling units, and patient cohorts remained fixed. Vertical dashed lines mark the 2% reference. **b**, Corresponding changes for individual FMs. Rows represent FMs, and columns represent sampling fractions. Blue and pink indicate negative and positive changes, respectively. White indicates zero change, including the 2% reference column. Color scales are determined separately for each heatmap. Bio-kNN denotes biological k-nearest-neighbor performance. CS, CI and RI denote clustering score, confounder insensitivity and robustness index, respectively. Balanced accuracy (BA) and average performance drop (APD) changes are expressed in percentage points (pp). Other changes retain their original metric units. Negative integrated Brier score (IBS) changes indicate lower prediction error. ID and OOD denote in-distribution and out-of-distribution evaluation. Mean tdAUC denotes mean time-dependent area under the receiver operating characteristic curve.

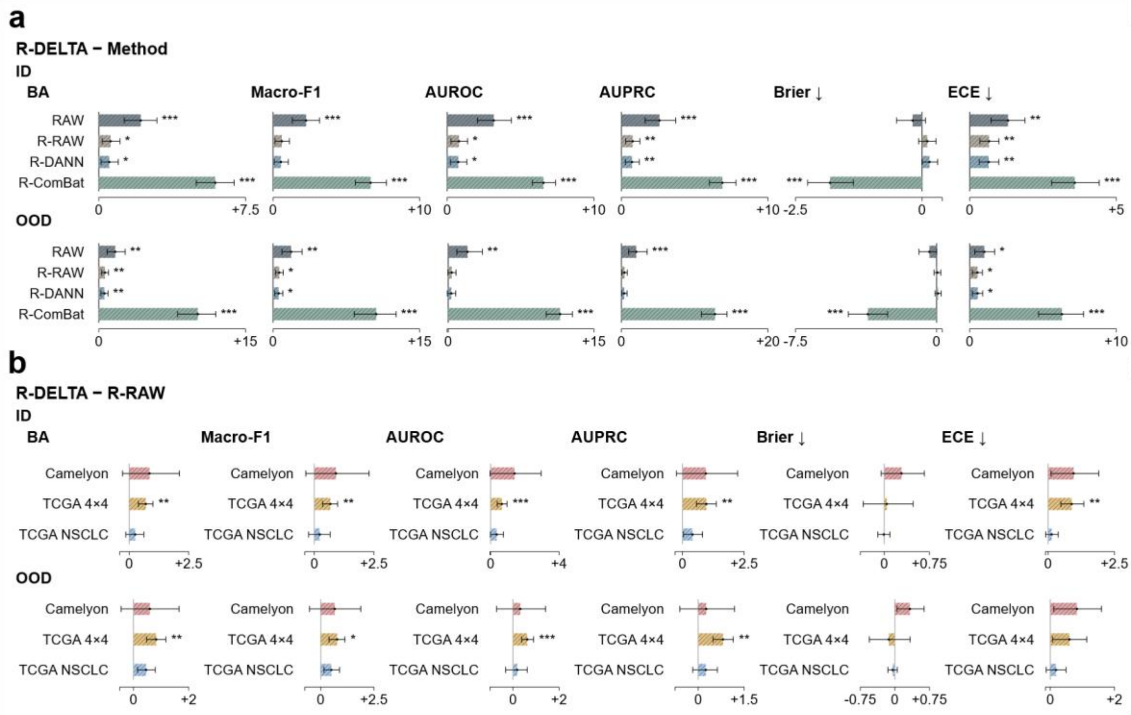


**c**

R-DELTA − R-RAW

| | BA | | Macro-F1 | | AUROC | | AUPRC | | Brier score ↓ | | ECE ↓ | |
|---|---|---|---|---|---|---|---|---|---|---|---|---|
| | ID | OOD | ID | OOD | ID | OOD | ID | OOD | ID | OOD | ID | OOD |
| BEPH | −0.01 | −0.36 | −0.07 | −0.49 | +0.53 | −0.03 | +0.47 | +0.21 | +0.14 | −0.19 | +0.22 | −0.18 |
| Ciga | +3.22 | +2.20 | +3.41 | +2.21 | +4.31 | +2.96 | +3.59 | +2.50 | −0.15 | −0.07 | +2.62 | +1.06 |
| CONCH | −0.42 | +1.58 | −0.47 | +1.58 | −0.67 | −0.07 | −0.50 | +0.25 | +0.24 | +0.07 | −0.40 | +1.59 |
| CONCH v1.5 | −0.95 | +0.19 | −0.93 | +0.03 | −0.65 | +0.35 | −0.44 | +0.40 | +0.49 | +0.27 | −0.52 | +0.68 |
| CTransPath | +0.36 | +0.24 | +0.51 | +0.34 | +0.80 | −1.18 | +0.76 | −1.05 | 0.00 | +0.03 | +0.51 | +0.14 |
| EXAONE-Path-2.5 | +0.61 | −0.22 | +0.59 | −0.29 | +0.40 | −0.19 | +0.55 | +0.09 | +0.16 | +0.26 | +0.99 | +0.11 |
| GPFM | +1.48 | +1.29 | +1.72 | +1.54 | +1.78 | +0.87 | +1.75 | +1.14 | −0.07 | +0.02 | +1.27 | +1.02 |
| H0-mini | +0.24 | −0.12 | +0.37 | +0.05 | +0.23 | −0.27 | +0.59 | +0.31 | −0.08 | +0.02 | +0.39 | +0.24 |
| H-optimus-0 | +0.19 | +0.67 | +0.25 | +0.68 | +0.64 | +0.42 | +0.55 | +0.35 | +0.11 | +0.12 | +0.36 | +0.83 |
| HIPT | +0.63 | +1.08 | +0.11 | +0.99 | +1.77 | +0.95 | +2.01 | +0.70 | −0.22 | −0.76 | +0.34 | −1.47 |
| Kaiko ViT-B/8 | −0.92 | −0.46 | −0.98 | −0.55 | −0.42 | −0.36 | −0.70 | −0.36 | +0.89 | +0.66 | +0.66 | +0.36 |
| MOOZY | +2.61 | +1.94 | +2.72 | +2.20 | +2.77 | +2.01 | +2.38 | +1.70 | −0.81 | −0.77 | +0.44 | +0.67 |
| MUSK | +1.10 | +1.07 | +1.22 | +1.33 | +0.63 | +0.56 | +0.56 | +0.37 | −0.15 | −0.05 | +0.91 | +0.88 |
| Original UNI | +0.60 | +0.25 | +0.60 | +0.20 | +1.27 | +0.19 | +0.96 | +0.23 | +0.64 | +0.55 | +0.67 | +1.28 |
| Phikon-v2 | +1.53 | +1.01 | +1.53 | +1.07 | +2.00 | +0.88 | +1.71 | +1.19 | +0.01 | +0.01 | +1.61 | +1.69 |
| Prov-GigaPath | +0.54 | −0.05 | +0.55 | −0.20 | +0.60 | +0.42 | +0.65 | +0.42 | +0.08 | +0.08 | +0.82 | +0.05 |
| RetCCL | −0.69 | −0.15 | −1.10 | −0.40 | −1.66 | −1.52 | −1.24 | −1.32 | +0.02 | +0.01 | −0.62 | −0.43 |
| UNI2-h | +0.70 | −0.09 | +0.79 | −0.09 | +0.88 | +0.28 | +0.72 | +0.10 | +0.06 | +0.08 | +0.72 | −0.20 |
| Virchow | +0.62 | +0.53 | +0.56 | +0.50 | +0.74 | +0.24 | +0.95 | +0.60 | +0.94 | +1.03 | +1.64 | +1.49 |
| Virchow2 | +0.58 | +1.87 | +0.66 | +2.17 | +0.12 | +0.96 | +0.10 | +0.98 | −0.16 | −0.38 | +0.58 | +0.88 |
| **Mean** | **+0.60*** | **+0.62**** | **+0.60** | **+0.64*** | **+0.80*** | **+0.37** | **+0.77**** | **+0.44** | **+0.11** | **+0.05** | **+0.66**** | **+0.53*** |
| Color scale | −3.82 / 0 / +3.82 | | −4.11 / 0 / +4.11 | | −6.03 / 0 / +6.03 | | −4.93 / 0 / +4.93 | | −1.20 / 0 / +1.20 | | −2.62 / 0 / +2.62 | |

**Supplementary Figure 2. Reinhard-DELTA in patch-aggregated cancer classification.**

Reinhard-DELTA (R-DELTA) was applied to embeddings from Reinhard-normalized patches before gated-attention aggregation. The same three tasks as in Fig. 3 were evaluated: tumor detection in Camelyon, four-class cancer classification in TCGA4×4, and lung adenocarcinoma versus squamous cell carcinoma classification in TCGA NSCLC. Balanced accuracy (BA) was the primary endpoint. Macro-F1, AUROC, AUPRC, Brier score and expected calibration error (ECE) provided complementary measures of discrimination and calibration. **a**, Mean paired differences between R-DELTA and the indicated comparators across 20 backbones. RAW denotes uncorrected embeddings from original images, R-RAW denotes uncorrected embeddings from Reinhard-normalized images, and R-DANN and R-ComBat denote the corresponding methods evaluated after Reinhard normalization. ComBat was evaluated transductively. **b**, Task-specific differences between R-DELTA and R-RAW. **c**, Corresponding differences for individual backbones. Each backbone estimate averages 100 fixed splits per task. All numerical effects are calculated as R-DELTA minus comparator and multiplied by 100. Positive differences favor R-DELTA for BA, macro-F1, AUROC and AUPRC, whereas negative differences favor R-DELTA for Brier score and ECE. ID and OOD denote in-distribution and out-of-distribution evaluation. In the heatmap, red indicates improvement and blue indicates deterioration, with the color mapping reversed for Brier score and ECE while numerical signs remain unchanged. Error bars show pointwise 95% confidence intervals from 10,000 bootstrap resamples of paired backbone effects. P values were obtained using two-sided exact paired signed-rank tests with Holm correction within comparison families. $^{*}P < 0.05$, $^{**}P < 0.01$ and $^{***}P < 0.001$.

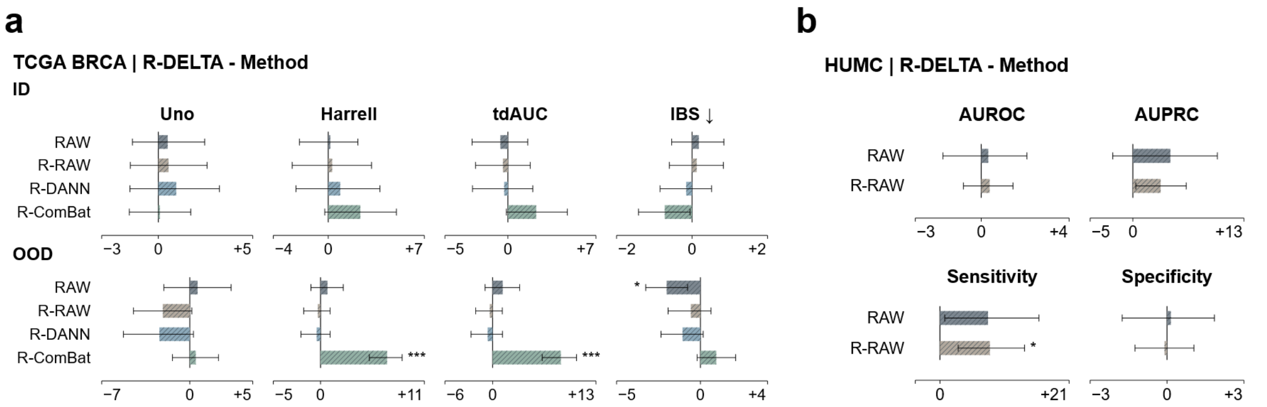


**c**

**TCGA BRCA | R-DELTA-R-RAW**

| | Uno | | Harrell | | tdAUC | | IBS ↓ | |
|---|---|---|---|---|---|---|---|---|
| | ID | OOD | ID | OOD | ID | OOD | ID | OOD |
| BEPH | +4.91 | +3.36 | +7.43 | +2.54 | +9.54 | +2.49 | −1.41 | +3.55 |
| Ciga | +4.15 | −2.22 | +1.68 | −3.31 | +0.84 | −4.22 | +0.28 | +0.00 |
| CONCH | −1.76 | −2.66 | +5.76 | −9.44 | +4.91 | −10.81 | −0.79 | −1.37 |
| CONCH v1.5 | +1.59 | −2.75 | −2.40 | +2.33 | −2.72 | +2.67 | −0.29 | −1.01 |
| CTransPath | −4.75 | −4.58 | −11.51 | −5.64 | −7.34 | −7.21 | −0.04 | +0.02 |
| EXAONE-Path-2.5 | −5.22 | −13.38 | −13.19 | +0.44 | −10.03 | +1.55 | +2.47 | +2.24 |
| GPFM | +1.30 | −6.24 | −2.64 | +1.65 | −3.67 | +2.42 | +0.29 | −0.50 |
| H-optimus-0 | +2.62 | +1.79 | +5.76 | −0.36 | +5.08 | +0.18 | −2.68 | −4.82 |
| H0-mini | +5.80 | −2.92 | −1.20 | +1.85 | −2.00 | +2.08 | +0.21 | +0.39 |
| HIPT | −0.66 | +4.48 | −1.68 | −1.18 | −0.39 | −2.12 | +0.20 | −2.31 |
| Kaiko ViT-B/8 | +8.05 | +5.97 | +1.20 | +1.20 | −0.23 | +1.08 | +0.00 | −0.37 |
| MOOZY | −1.97 | +6.61 | +3.84 | −0.39 | +0.70 | −2.35 | +0.29 | −3.63 |
| MUSK | −2.14 | −3.92 | +6.00 | −4.62 | +2.66 | −4.36 | +1.60 | −1.92 |
| Original UNI | −3.21 | −1.40 | −3.60 | +2.28 | −5.77 | +3.19 | +0.53 | +1.42 |
| Phikon-v2 | −3.38 | −9.64 | +2.64 | +0.13 | +0.60 | +1.23 | −1.09 | −2.74 |
| Prov-GigaPath | −0.48 | +1.18 | +15.59 | +3.05 | +6.48 | +4.19 | −3.93 | −8.91 |
| RetCCL | +6.63 | −7.88 | +2.88 | −1.31 | +2.93 | −3.00 | +0.10 | +2.63 |
| UNI2-h | −9.47 | −4.69 | −6.71 | −0.23 | −5.30 | +0.00 | +3.37 | +2.28 |
| Virchow | +1.26 | −8.19 | −6.95 | +3.82 | −7.03 | +3.43 | +2.26 | +2.71 |
| Virchow2 | +7.59 | +4.13 | +2.88 | +1.37 | +2.83 | +2.39 | +1.09 | +0.86 |
| **Mean** | **+0.54** | **−2.15** | **+0.29** | **−0.29** | **−0.40** | **−0.36** | **+0.12** | **−0.57** |

Color scales: -13.4 0 +8.1; -13.2 0 +15.6; -10.8 0 +9.5; -8.9 0 +3.6

**d**

**HUMC | R-DELTA-R-RAW**

| | AUROC | AUPRC | Sens. | Spec. |
|---|---|---|---|---|
| BEPH | −0.53 | +0.28 | +9.38 | −2.00 |
| Ciga | +2.12 | +9.79 | +12.50 | +0.00 |
| CONCH | +2.16 | +12.05 | +21.88 | +0.00 |
| CONCH v1.5 | +0.19 | +0.20 | +3.12 | −1.00 |
| CTransPath | −0.66 | −0.30 | −3.12 | +0.00 |
| EXAONE-Path-2.5 | −0.16 | +0.92 | +3.12 | +0.00 |
| GPFM | +0.28 | +5.27 | +9.38 | +1.00 |
| H-optimus-0 | +0.53 | +1.57 | +9.38 | +0.00 |
| H0-mini | −1.06 | −0.82 | −3.12 | +1.00 |
| HIPT | +0.38 | +9.63 | +15.62 | +1.00 |
| Kaiko ViT-B/8 | +0.00 | −0.44 | +6.25 | +1.00 |
| MOOZY | +1.41 | +10.97 | +21.88 | −1.00 |
| MUSK | +1.72 | +3.13 | +15.62 | −1.00 |
| Original UNI | +0.56 | +1.87 | +9.38 | −2.00 |
| Phikon-v2 | +0.50 | +4.41 | +6.25 | +2.00 |
| Prov-GigaPath | +0.34 | +0.97 | −9.38 | +0.00 |
| RetCCL | −0.44 | +3.38 | +18.75 | −1.00 |
| UNI2-h | −1.66 | −1.09 | −6.25 | −1.00 |
| Virchow | +1.53 | +5.05 | +12.50 | +0.00 |
| Virchow2 | +0.44 | −1.73 | +9.38 | +1.00 |
| **Mean** | **+0.38** | **+3.25** | **+8.12*** | **−0.10** |

Color scales: -1.7 +2.2; -1.7 +12.1; -9.4 +21.9; -2.0 +2.0

**Supplementary Figure 3. WSI-level predictive performance of DELTA after Reinhard stain normalization.**

R-DELTA denotes DELTA applied to patch embeddings extracted from Reinhard-normalized images before downstream prediction. **a,c**, TCGA-BRCA survival prediction under in-distribution (ID) and held-out-center out-of-distribution (OOD) evaluation. Uno C-index was the primary endpoint, complemented by Harrell C-index, mean time-dependent AUC (tdAUC) and integrated Brier score (IBS). **b,d**, HUMC normative cancer screening, assessed using AUROC, AUPRC, sensitivity and specificity from pooled out-of-fold case predictions. Sensitivity and specificity were evaluated at training-derived thresholds. **a,b**, Mean paired differences across 20 backbones. TCGA-BRCA comparators include RAW, R-RAW, R-DANN and R-ComBat. RAW denotes uncorrected embeddings from original images. R-RAW, R-DANN and R-ComBat denote uncorrected embeddings, domain adversarial neural networks and transductive ComBat, respectively, following Reinhard stain normalization. HUMC comparators include RAW and R-RAW. **c,d**, Corresponding differences between R-DELTA and R-RAW for individual backbones. All numerical effects are calculated as R-DELTA minus comparator and multiplied by 100. Positive differences favor R-DELTA except for IBS, for which negative differences favor R-DELTA. Heatmap colors indicate improvement (red) and deterioration (blue), with the color mapping reversed for IBS while numerical signs remain unchanged. Error bars show pointwise 95% confidence intervals from 20,000 bootstrap resamples. TCGA-BRCA intervals resample paired backbone effects. Two-sided exact paired sign-flip P values were Holm-adjusted across the four prespecified comparators within each endpoint and evaluation domain in the Reinhard input branch. HUMC intervals use hierarchical paired resampling of cases within cancer-status strata and backbones. Two-sided centered hierarchical bootstrap P values were Holm-adjusted across the same five prespecified DELTA and R-DELTA contrasts within each metric used in Figure 4. Heatmap significance symbols refer to mean paired differences and use the same adjusted P values as the corresponding bar plots. *P < 0.05, **P < 0.01 and ***P < 0.001.

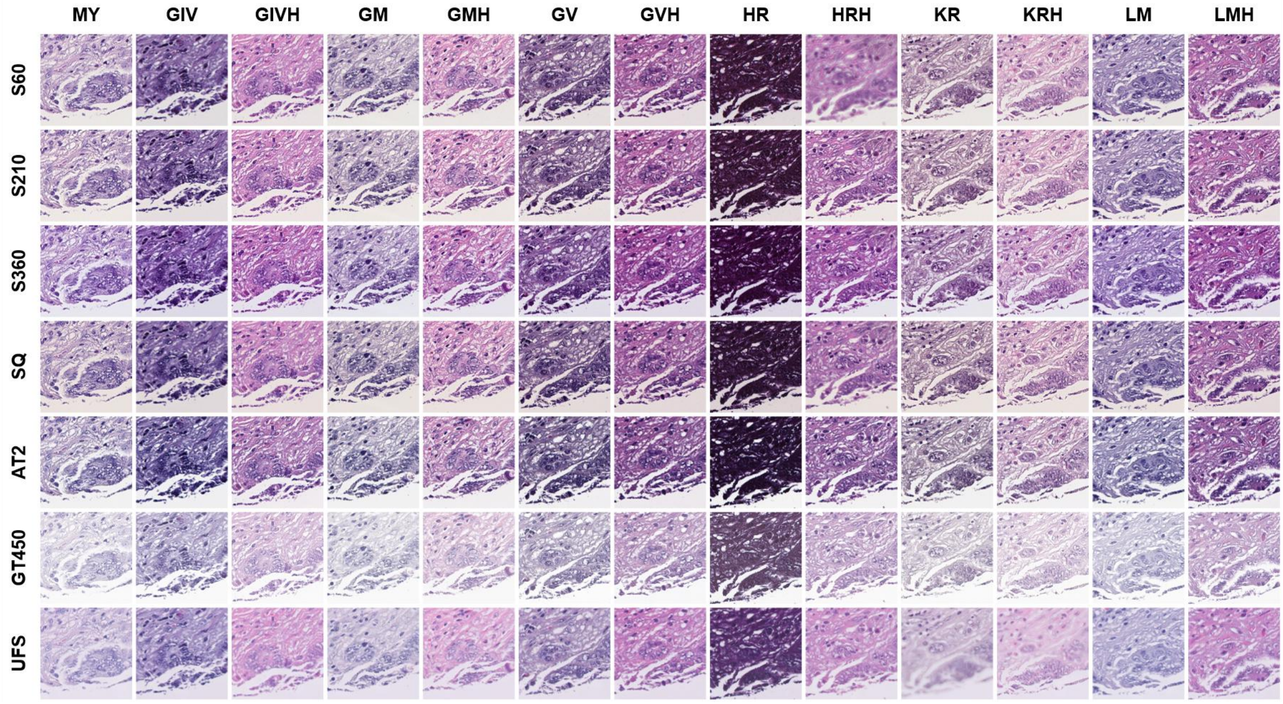


**Supplementary Figure 4. Scanner and staining-condition variation in PLISM.**

A representative registered metastatic colorectal cancer region is shown across 13 H&E staining conditions and seven whole-slide scanners. Rows represent scanners and columns represent staining protocols. All 91 panels show uncorrected input images. Cross-stain images originate from registered serial sections and therefore include section-related tissue differences in addition to staining variation. H&E: hematoxylin and eosin. MY: Mayer. GIV and GIVH: Gill IV. GM: New Type G. GMH: GM hematoxylin. GV and GVH: Gill V. HR and HRH: Harris. KR and KRH: Carrazi. LM and LMH: Lillie–Mayer. H suffix: protocols selected for greater color separation between the hematoxylin and eosin components. S60: Hamamatsu NanoZoomer S60. S210: Hamamatsu NanoZoomer S210. S360: Hamamatsu NanoZoomer S360. SQ: Hamamatsu NanoZoomer SQ. AT2: Leica Aperio AT2. GT450: Leica Aperio GT450. UFS: Philips Ultrafast Scanner.

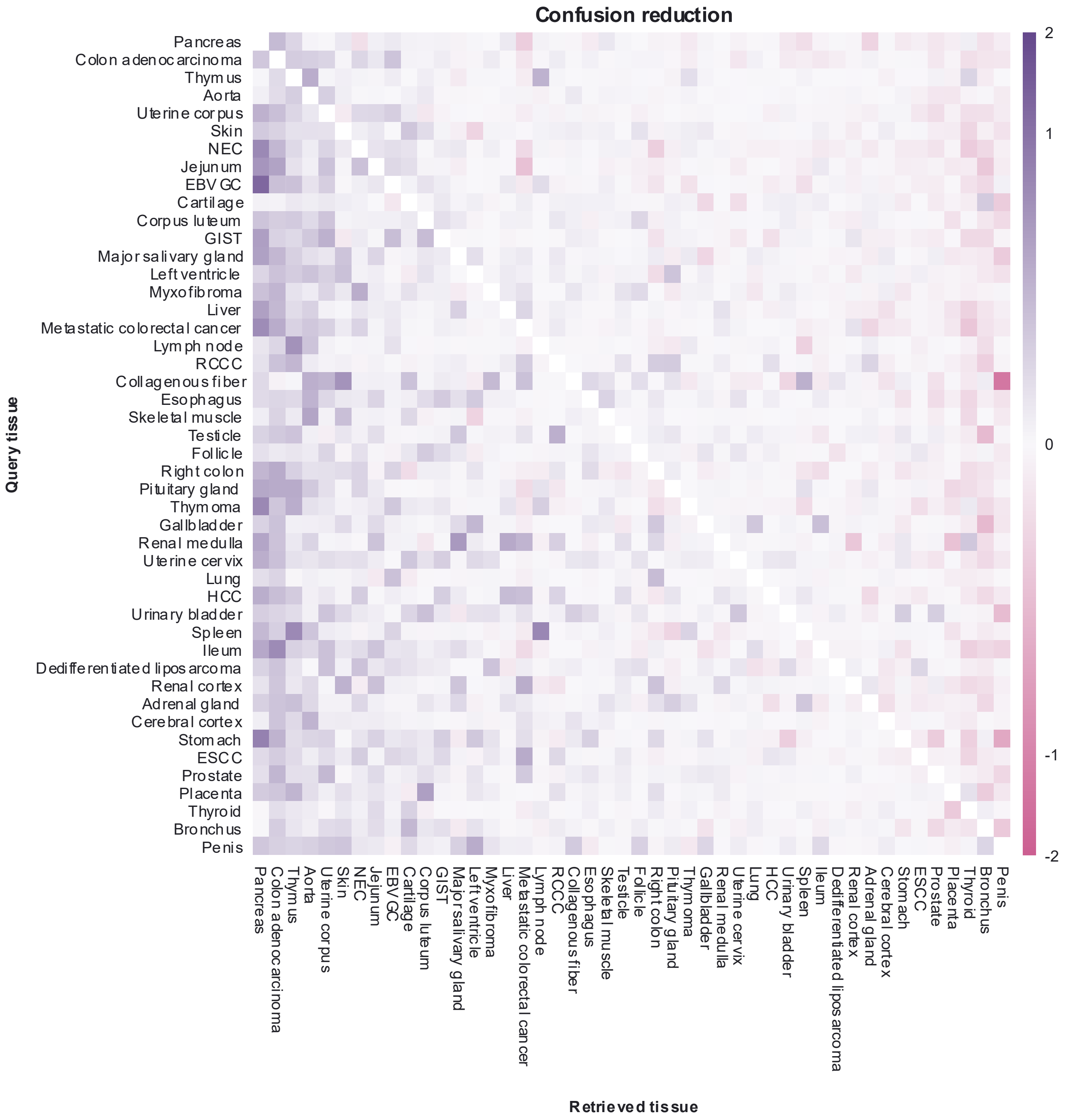


**Supplementary Figure 5. DELTA-associated changes in tissue-retrieval confusion in PLISM.**

Rows represent 46 query tissue groups and columns represent the tissue labels of the nearest retrieved regions. Tissue annotations include NEC (neuroendocrine carcinoma), EBVGC (Epstein–Barr virus-associated gastric cancer), GIST (gastrointestinal stromal tumor), RCCC (renal clear cell carcinoma), HCC (hepatocellular carcinoma) and ESCC (esophageal squamous cell carcinoma). The corresponding registered region was excluded from the reference gallery for each query. Each off-diagonal cell shows the difference between RAW (uncorrected embeddings) and DELTA in row-normalized retrieval frequency, calculated as RAW minus DELTA and expressed in percentage points. Results were averaged equally across 20 backbones and three acquisition-change types: scanner, stain-condition and combined changes. Purple indicates reduced cross-tissue confusion and pink indicates increased confusion. Both axes use an identical tissue order, ranked by decreasing mean off-diagonal confusion reduction within each retrieved-tissue column across the other 45 query groups. Diagonal cells are omitted. A shared inverse-hyperbolic-sine color transformation is applied, with color-bar labels retaining the original percentage-point units. Cell values are descriptive summaries without cell-wise significance testing.

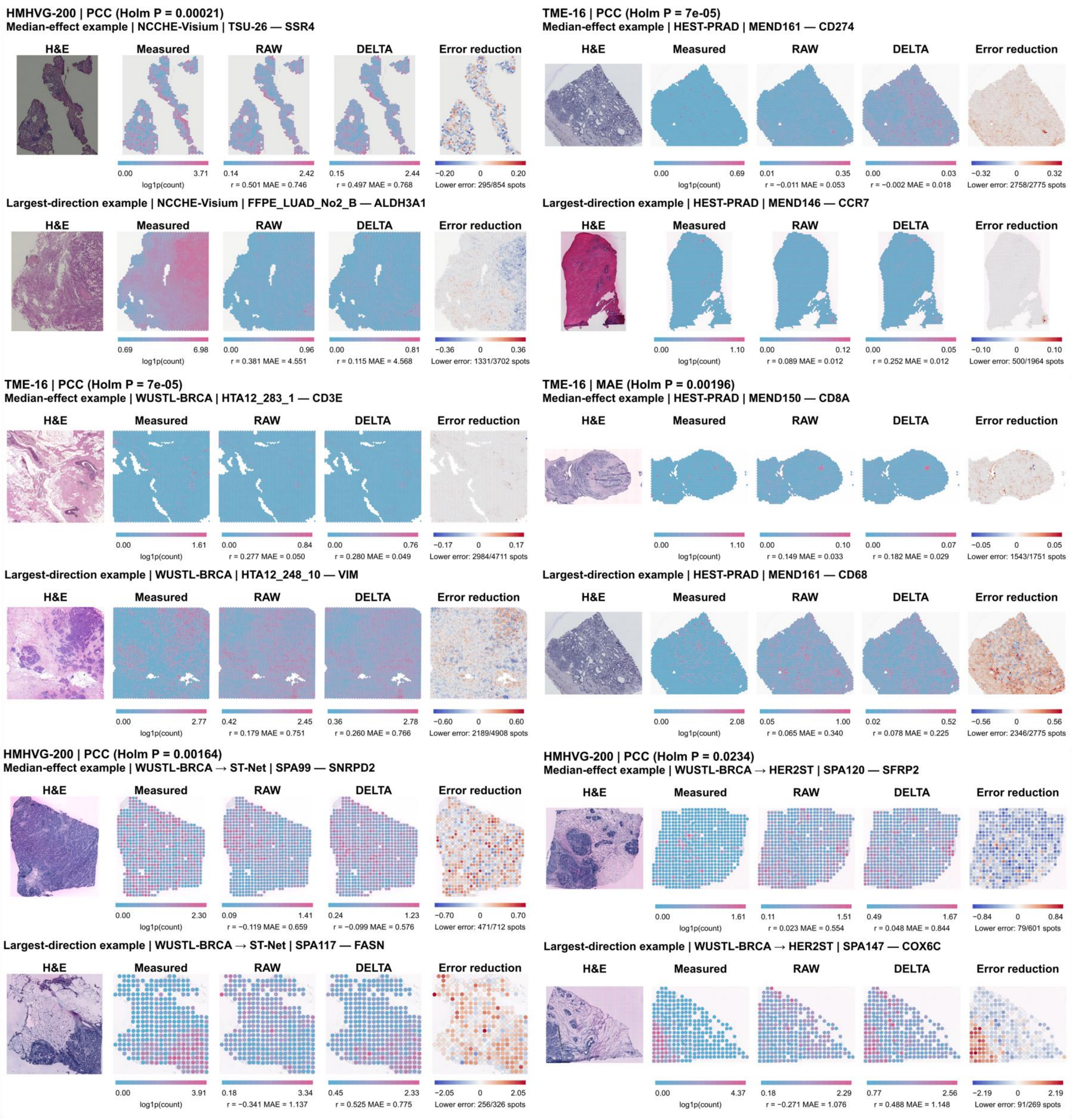


**Supplementary Figure 6. Spatial examples of gene-expression prediction after DELTA correction.**

Measured expression is compared with predictions from raw and DELTA-corrected UNI-v2 embeddings. Each example represents one gene within a tissue section from a comparison with Holm-adjusted P < 0.05. For each comparison, the median-effect example is the eligible section–gene pair nearest the median change in the indicated endpoint. The largest-direction example has the greatest change in the direction of the dataset-level result among pairs from another tissue section. Displayed P values refer to dataset or external-transfer comparisons. Sections formed the paired units within fixed folds in the original benchmark, and patients formed the paired units for ST-Net and HER2ST. Two-sided sign-flip P values were Holm-adjusted across tasks separately within each evaluation group, gene panel and endpoint. Expression maps display log1p-transformed counts with separate color limits. Blue and pink indicate lower and higher expression, respectively. Error reduction is RAW absolute error minus DELTA absolute error at each spot. Red and blue indicate lower and higher error after correction, respectively, on scales symmetric about zero. Counts report spots with lower DELTA error relative to all evaluated spots. Displayed Pearson correlation coefficients (r) and mean absolute errors (MAE) were calculated across all spots for the indicated gene and section.

### Supplementary Tables

**Supplementary Table 1. Absolute patch-level representation and classification performance.**

| Endpoint | RAW | R-RAW | DANN | R-DANN | ComBat | R-ComBat | DELTA | R-DELTA |
|---|---|---|---|---|---|---|---|---|
| Bio-kNN | 0.8807 | 0.8789 | 0.8854 | 0.8837 | 0.8917 | 0.8885 | 0.8848 | 0.8828 |
| CS | 0.3481 | 0.4173 | 0.4014 | 0.4412 | 0.4655 | 0.4664 | 0.4603 | 0.5104 |
| CI | 0.5621 | 0.6625 | 0.6088 | 0.7262 | 1.1162 | 1.1685 | 0.6630 | 0.7150 |
| RI | 0.6185 | 0.7259 | 0.6643 | 0.7645 | 0.8104 | 0.8391 | 0.7165 | 0.7517 |
| ID BA | 0.8455 | 0.8535 | 0.8447 | 0.8533 | 0.8707 | 0.8711 | 0.8571 | 0.8584 |
| ID APD | -0.0529 | -0.0425 | -0.0531 | -0.0417 | -0.0219 | -0.0212 | -0.0380 | -0.0358 |
| OOD BA | 0.8361 | 0.8472 | 0.8329 | 0.8467 | 0.8101 | 0.8153 | 0.8456 | 0.8513 |
| OOD APD | -0.0345 | -0.0191 | -0.0377 | -0.0185 | -0.0726 | -0.0649 | -0.0212 | -0.0129 |

Values average Camelyon, TCGA4x4 and Tolkach ESCA within each encoder and then the 20-encoder panel, retaining the original metric scales. Bio-kNN denotes biological nearest-neighbor performance, CS clustering score, CI confounder insensitivity and RI robustness index. BA denotes balanced accuracy. APD retains its signed native scale relative to the common RAW balanced-condition reference and is not expressed in percentage points here. ID and OOD denote in-distribution and out-of-distribution evaluation. RAW denotes uncorrected embeddings and R denotes Reinhard-normalized input. ComBat was evaluated transductively. Full paired comparisons, confidence intervals and adjusted P values are provided in Source Data.

**Supplementary Table 2. Absolute WSI-level survival performance in TCGA-BRCA.**

| Endpoint | RAW | R-RAW | DANN | R-DANN | ComBat | R-ComBat | DELTA | R-DELTA |
|---|---|---|---|---|---|---|---|---|
| ID Uno C-index | 0.6104 | 0.6099 | 0.5986 | 0.6058 | 0.6145 | 0.6145 | 0.6442 | 0.6153 |
| ID Harrell C-index | 0.6777 | 0.6764 | 0.6652 | 0.6705 | 0.6594 | 0.6559 | 0.7112 | 0.6793 |
| ID mean tdAUC | 0.7308 | 0.7288 | 0.7254 | 0.7278 | 0.7086 | 0.7023 | 0.7600 | 0.7248 |
| ID IBS | 0.0999 | 0.1004 | 0.1028 | 0.1032 | 0.1120 | 0.1089 | 0.0990 | 0.1016 |
| OOD Uno C-index | 0.4717 | 0.4993 | 0.4656 | 0.5020 | 0.4521 | 0.4731 | 0.4730 | 0.4778 |
| OOD Harrell C-index | 0.5683 | 0.5786 | 0.5683 | 0.5797 | 0.5037 | 0.5051 | 0.5714 | 0.5757 |
| OOD mean tdAUC | 0.5914 | 0.6077 | 0.5929 | 0.6103 | 0.5194 | 0.5182 | 0.5978 | 0.6040 |
| OOD IBS | 0.1327 | 0.1183 | 0.1415 | 0.1232 | 0.1067 | 0.1031 | 0.1221 | 0.1125 |

Values are means across 20 encoders on fixed patient partitions. Patient risk predictions were averaged across three optimization seeds before metric calculation. Uno C-index was the primary endpoint. Harrell C-index and mean time-dependent AUC provide complementary discrimination measures. IBS denotes integrated Brier score, for which lower values indicate lower error. ID and OOD test sets contain 80 and 451 patients, respectively. RAW denotes uncorrected embeddings and R denotes Reinhard-normalized input. Full paired comparisons and model-panel uncertainty estimates are provided in Source Data.

**Supplementary Table 3. Absolute performance in HUMC normative screening.**

| Representation | AUROC | AUPRC | Sensitivity | Specificity |
|---|---|---|---|---|
| RAW | 0.9551 | 0.8304 | 0.7563 | 0.9415 |
| R-RAW | 0.9545 | 0.8418 | 0.7531 | 0.9440 |
| DELTA | 0.9620 | 0.8786 | 0.8563 | 0.9425 |
| R-DELTA | 0.9583 | 0.8743 | 0.8344 | 0.9430 |

Values are means of encoder-specific pooled out-of-fold metrics across 20 encoders, rather than predictions from an ensemble. The primary analysis retained five Lloyd iterations for bank construction. The cohort comprises 100 non-tumor and 32 cancer cases from distinct patients. Sensitivity and specificity were measured at branch-specific thresholds determined from the 95th percentile of leave-one-case-out training non-tumor scores. R-RAW and R-DELTA denote the corresponding Reinhard input branches. Hierarchical case-plus-encoder confidence intervals and the complete prespecified comparison family are provided in Source Data.

**Supplementary Table 4. Distance reduction and tissue retrieval across seven PLISM scanners under MY staining.**

| Method | Squared embedding-distance reduction | Matched-field top-1 retrieval | Within-tissue top-1 | Reciprocal rank | Tissue-category concordance |
|---|---|---|---|---|---|
| Raw encoder embeddings | 0.0000 | 0.8273 | 0.8476 | 0.8655 | 0.8228 |
| **DELTA** | **0.1923** | **0.8492** | **0.8670** | **0.8842** | **0.8300** |
| Fitting-bank PCA | 0.1277 | 0.8242 | 0.8455 | 0.8621 | 0.8191 |
| Unpaired displacement | 0.1503 | 0.8286 | 0.8493 | 0.8662 | 0.8202 |
| Random subspace | 0.0179 | 0.8272 | 0.8475 | 0.8654 | 0.8227 |

Values average tissue-category results within encoder and then the 20-encoder panel. The PLISM MY-staining subset contains 3,417 corresponding tissue fields on seven scanners, yielding 21 scanner pairs across 46 tissue categories. All operators were estimated from TCGA-BRCA and transferred without PLISM-specific estimation. PCA, unpaired displacement and random-subspace controls share DELTA's encoder-specific rank and center. Squared embedding-distance reduction is the fraction of original squared Euclidean distance removed by projection and is zero for raw encoder embeddings. Each non-S60 scanner is a query source, with S60 as the fixed retrieval gallery. Random-subspace results average metrics from five frozen bases. The MY-only analysis is descriptive. Figure 5 and Supplementary Data 1 and Source Data report the expanded analysis across 13 stain conditions.

**Supplementary Table 5. Software configurations of the principal execution environments.**

| Software | Python 3.9 environment | Python 3.10 environment |
|---|---|---|
| Python | 3.9.23 | 3.10.18 |
| PyTorch | 2.1.0 | 2.8.0 |
| torchvision | 0.16.0 | 0.23.0 |
| scikit-learn | 1.5.2 | 1.7.2 |
| SciPy | 1.13.1 | 1.15.3 |
| CUDA for the PyTorch build | 12.1 | 12.8 |

The CUDA versions identify the corresponding PyTorch builds. Assessment-specific environments are described in Supplementary Methods S18.
